# The distribution of onion virulence gene clusters among *Pantoea* spp

**DOI:** 10.1101/2020.12.18.423501

**Authors:** Shaun P. Stice, Gi Yoon Shin, Stefanie De Armas, Santosh Koirala, Guillermo A. Galván, María Inés Siri, Paul M. Severns, Teresa Coutinho, Bhabesh Dutta, Brian H. Kvitko

**Affiliations:** College of Agriculture and Environmental Science, Department of Plant Pathology, University of Georgia, Athens, Georgia, United States of America; Centre for Microbial Ecology and Genomics, Forestry and Agriculture Biotechnology Institute, Department of Biochemistry Genetics and Microbiology, University of Pretoria, Pretoria, South Africa; Facultad de Química, Área Microbiología, Departamento de Biociencias, Universidad de la República, Montevideo, Uruguay; Facultad de Agronomía, Centro Regional Sur (CRS), Departamento de Producción Vegetal, Universidad de la República, Canelones, Uruguay

**Keywords:** Pantoea ananatis, Pantoea spp., onion, virulence gene, bulb rotting, gene cluster distrubution

## Abstract

*Pantoea ananatis* is a gram-negative bacterium and the primary causal agent of center rot of onions in Georgia. Previous genomic studies identified two virulence gene clusters, HiVir and *alt*, associated with center rot. The HiVir gene cluster is required to induce necrosis on onion tissues via synthesis of a predicted small molecule toxin. The *alt* gene cluster aids in tolerance to thiosulfinates generated during onion tissue damage. Whole genome sequencing of other *Pantoea* species suggest that these gene clusters are present outside of *P. ananatis*. To assess the distribution of these gene clusters, two PCR primer sets were designed to detect the presence of HiVir and *alt*. Two hundred fifty-two strains of *Pantoea* spp. were phenotyped using the red onion scale necrosis (RSN) assay and were assayed using PCR for the presence of these virulence genes. A diverse panel of strains from three distinct culture collections comprised of 24 *Pantoea* species, 41 isolation sources, and 23 countries, collected from 1946-2019, were tested. There is a significant association between the *alt* PCR assay and *Pantoea* strains recovered from symptomatic onion (*P*<0.001). There is also a significant association of a positive HiVir PCR and RSN assay among *P. ananatis* strains but not among *Pantoea* spp., congeners. This may indicate a divergent HiVir cluster or different pathogenicity and virulence mechanisms. Last, we describe natural *alt* positive [RSN^+^/HiVir^+^/*alt*^*+*^] *P. ananatis* strains, which cause extensive bulb necrosis in a neck-to-bulb infection assay compared to *alt* negative [RSN^+^/HiVir^+^/*alt*^*-*^] *P. ananatis* strains. A combination of assays that include PCR of virulence genes [HiVir and *alt*] and an RSN assay can potentially aid in identification of onion-bulb-rotting pathogenic *P. ananatis* strains.

## 1 Introduction

Onion center rot is an economically impactful disease that routinely results in significant losses to the yield and marketability of onion (*Allium cepa* L.). Symptoms of the disease include necrotized leaves and rotted bulbs (Gitaitis and Gay, 1997). The causal agent of onion center rot is primarily *Pantoea ananatis* in the southeastern United States; however, bacterial onion blights and bulb rots caused by other *Pantoea* species, including *Pantoea allii, P. agglomerans, P. stewartii* subsp. *indologenes*, and *P. dispersa*, have been reported in the literature (Edens et al., 2006; Brady et al., 2011; Chang et al., 2018; Stumpf et al., 2018). *P. ananatis* is transmitted by thrips and can move from infected onion leaves to the corresponding onion scales (Carr et al., 2013; Dutta et al., 2016). Recent comparative and functional genetic studies have determined that the HiVir and *alt* gene clusters function together to drive necrotrophic infection of onion by virulent *P. ananatis* strains (Stice et al., 2020).

The HiVir gene cluster is a chromosomal gene cluster in *P. ananatis* that is hypothesized to code for the synthesis of a yet undescribed phosphonate secondary metabolite that acts as a plant toxin (Asselin et al., 2018; Takikawa and Kubota, 2018). Deletion of the first gene in the HiVir cluster *pepM*, a phosphoenolpyruvate mutase gene, renders strains unable to induce necrotic lesions on onion foliage and tobacco leaves. The mutant strains were also unable to induce necrosis on detached red onion scales after three days of inoculation (red onion scale necrosis, RSN), compared to the wild-type (WT) strain (Asselin et al., 2018; Stice et al., 2020). The HiVir cluster, which was originally identified as a potential insecticidal toxin cluster by De Maayer *et al*. 2014, is common to several plant pathogenic *P. ananatis* strains (De Maayer et al., 2014). The HiVir cluster is present in the genomes of the *P. ananatis* type strain LMG 2665^T^ (GenBank: JMJJ00000000; EL29_RS14675-RS14595) and the strain LMG 20103 (GenBank: NC_013956; PANA_RS16700-RS16615), which are the causal agents of pineapple fruitlet rot and Eucalyptus blight, respectively (De Maayer et al., 2010; Adam et al., 2014). *P. ananatis* strains reported as non-pathogenic on onion lacked the HiVir cluster (Asselin et al., 2018).

The plasmid-borne gene cluster *alt* encodes a cohort of enzymes that confer tolerance to the antimicrobial thiosulfinates released by damaged *Allium* tissues (Stice et al., 2020). We previously characterized the function of the *alt* cluster in *P. ananatis* by creating autobioluminescently-labeled PNA 97-1R isogenic mutants (Δ*alt*, Δ*pepM*, and Δ*alt*Δ*pepM*) and by testing them using the RSN assay and a neck-to-bulb infection/colonization assay. The later assay involves inoculating onion plants (bulb-swelling stage) at the neck with a bacteria-soaked toothpick and evaluating them at harvest maturity for *P. ananatis* colonization using long exposure imaging to determine bacterial colonization patterns. The strain PNA 97-1R WT [HiVir^+^/*alt*^+^] was RSN^+^ and caused a bulb rot in the neck-to-bulb infection assay, with extensive colonization supported by a bioluminescent signal throughout the symptomatic tissue. The strain PNA 97-1R Δ*alt* [HiVir^+^/*alt*^-^] was also RSN^+^ but did not cause a bulb rot in the neck-to-bulb infection assay and lacked a bioluminescent signal in the tissue. The strain PNA 97-1R Δ*pepM* [HiVir^-^/*alt*^+^] and Δ*alt*Δ*pepM* [HiVir^-^/*alt*^-^] were RSN^-^, did not cause a bulb rot, and did not produce bioluminescent signal in the neck-to-bulb assay (Stice et al., 2020). The qualitative difference of bulb rot symptoms in the neck-to-bulb infection assay between the WT and the Δ*alt* mutant strains indicated that the presence or absence of this cluster is important for bulb colonization and center rot symptoms in onion (Stice et al., 2020). Considering that the HiVir cluster is required for foliar necrosis in various *P. ananatis* strains and both the HiVir and *alt* clusters are required for the neck-to-bulb colonization of the onion bulb, we hypothesize that aggressive bulb rotting strains should contain both the *alt* and the HiVir clusters.

Though the HiVir and *alt* clusters have been functionally studied in *P. ananatis*, these clusters are distributed among other *Pantoea* spp. that have been reported to cause onion center rot disease (Stumpf et al., 2018; Stice et al., 2020). Using BlastX search schemes in the NCBI GenBank Database, we identified homologous HiVir clusters in several strains of *P. agglomerans* and *P. stewartii* subsp. *indologenes* and homologous *alt* clusters in several strains of *P. agglomerans, P. stewartii* subsp. *indologenes*, and *P. allii*. Hence, it would be useful to assess the distribution of both gene clusters amongst *Pantoea* spp. isolated from onion and non-onion sources. Here we describe two PCR assays to detect the pathogenicity and virulence factors (HiVir and the *alt* virulence genes) and the RSN assay for determining the onion pathogenic potential of *Pantoea* spp. strains. We utilized these three assays to analyze three distinct culture collections (*n*=252 strains) comprised of *Pantoea* spp. strains of environmental, pathogenic, epiphytic, and endophytic origins. In addition, we characterized three natural *alt* positive [RSN^+^/HiVir^+^/*alt*^*+*^] and three natural *alt* negative [RSN^+^/HiVir^+^/*alt*^*-*^] *P. ananatis* strains based on the symptomatology of onions when using a neck-to-bulb infection assay.

## 2 Materials and Methods

### 2.1 Bacterial strains and culture conditions

A total of 252 *Pantoea* spp. strains were selected from three independently curated culture collections: University of Georgia – Coastal Plain Experiment Station (UGA-CPES) (*n*=74 strains), University of Pretoria – Bacterial Culture Collection (UP-BCC) (*n*=124 strains), and Universidad de la República – Microorganisms of Agricultural Importance (UR-MAI) (*n*=54 strains). All bacterial strains were stored under cryopreservation at their respective institutions at −80 °C. Strains were routinely cultured from single clones recovered on either nutrient agar (NA) or lysogeny broth (LB) agar parent plates and were grown in 5 mL of NA/LB liquid media in 14 mL glass culture tubes at 28°C with shaking.

### 2.2 Description of *Pantoea* culture collections

#### 2.2.1 Strain selection

Each institution selected strains from their respective collections to include diverse *Pantoea* spp. and isolation sources, with specific enrichment of *P. ananatis* strains and *Pantoea* spp. strains isolated from diseased onion. Biases in strain selection were unavoidable; however, we employed relevant statistical methods to account for biases in our data sets.

#### 2.2.2 University of Georgia Coastal Plains Experiment Station (UGA-CPES)

Strains selected from the University of Georgia Coastal Plains Experiment Station (UGA-CPES) consisted of 4 *Pantoea* species isolated between 1983 and 2018: *P. ananatis* (*n*=69), *P. agglomerans* (*n*=2), *P. allii* (*n*=1), and *P. stewartii* subsp. *indologenes* (*n*=2). UGA-CPES-selected strains were primarily isolated from onion bulbs/leaves exhibiting center rot symptoms (*n*=43) and onion seeds (*n*=2), whereas the remainder of the strains were isolated from environmental sources, either from weeds (*n*=17) or thrips (*n*=12), using semi-selective PA20 agar (Goszczynska et al., 2006).

#### 2.2.3 University of Pretoria Bacterial Culture Collection (UP-BCC)

Strains selected from the University of Pretoria Bacterial Culture Collection (UP-BCC) consisted of 17 *Pantoea* species isolated between 1946 and 2015: *P. agglomerans* (*n*=18), *P. allii* (*n*=16), *P. ananatis* (*n*=44), *P. anthophila* (*n*=5), *P. bejingensis* (*n*=2), *P. conspicua* (*n*=1), *P. deleyi* (*n*=1), *P. dispersa* (*n*=6), *P. eucalypti* (*n*=6), *P. eucrina* (*n*=3), *P. pleuroti* (*n*=2), *P. rodasii* (*n*=1), *P. rwandensis* (*n*=1), *P. stewartii* (*n*=10), *P. vagans* (*n*=4), *P. wallisii* (*n*=2), and *P*. spp. [species not determined] (*n*=2). UP-BCC-selected strains were primarily isolated from onion (*n*=33), Eucalyptus (*n*=23), and maize (*n*=22). Seventy-eight of the UP-BCC strains were isolated from other sources ranging from human wounds to watermelon. Thirty-one of the selected strains in the UP-BCC are from the Belgian Coordinated Collections of Microorganisms (BCCM/LMG Bacteria Culture Collection).

#### 2.2.4 Universidad de la República Microorganisms of Agricultural Importance (UR-MAI)

Strains selected from the Universidad de la República Microorganisms of Agricultural Importance (UR-MAI) collection consisted of six *Pantoea* spp. isolated between 2015 and 2019: *P. agglomerans* (*n*=11), *P. allii* (*n*=4), *P. ananatis* (*n*=2), *P. eucalypti* (*n*=33), *P. vagans* (*n*=1), and *P*. spp. [species not determined] (*n*=3). UR-MAI-selected strains were isolated from streaked and spotted onion leaves, seed-stalks, and decayed onion bulbs.

### 2.3 DNA extraction

#### 2.3.1 UGA-CPES

Bacterial genomic DNA was extracted from overnight (O/N) cultures using the Gentra Puregene Yeast/Bact Kit (Qiagen) (25/74 strains). The purified DNA was adjusted to a final concentration of 80-50 ng/µL with Tris-EDTA buffer (10mM Tris-HCL, 1 mM EDTA, pH 8.0) and stored at 4 °C until use. DNA was also extracted using a DNA boil-prep protocol (49/74 strains). A single colony was selected from a culture plate, removed with a sterile toothpick, and agitated in a 200 µL PCR tube with 50 µL of sterile nuclease-free water. The PCR tube was heated to 95 °C for 20 m with a hold at 4 °C. Boil-preps were stored at 4 °C until use.

#### 2.3.2 UP-BCC

Bacterial genomic DNA was extracted using the Zymo Quick-DNA miniprep kit (Zymo Research) or PrepMan extraction kit (Applied Biosystems) according to the manufacturer instructions.

#### 2.3.3 UR-MAI

Bacterial suspensions were grown O/N in NBY after transferring single colonies of each strain from 48 h cultures on NBY-agar medium on a rotary shaker at 150 rpm. Thereafter, 2 mL of bacterial suspensions were used for DNA extraction following the protocol described by Ausubel *et al*. 2003 (Ausubel et al., 2003). Final DNA concentrations were adjusted to 50 ng/µL and stored until use.

### 2.4 HiVir PCR assay design

The HiVir2p_F/R primer pair was designed to detect *P. ananatis* strains with the ability to induce the red scale necrosis (RSN) phenotype associated with the HiVir cluster (Stice et al., 2020). HiVir2p_F/R was designed using a MAUVE alignment of the HiVir cluster and the Primer3 (v2.3.7) plugin within Geneious Prime (v 2019.1.3) (Darling et al., 2004). The HiVir clusters of PNA 97-1R (GenBank: CP020943.2; B9Q16_20825-20770) and LMG 2665^T^ (GenBank: NZ_JFZU01000015; CM04_RS24645-RS25770) [RSN^+^/HiVir^+^] were aligned with three genome-sequenced strains carrying a HiVir cluster that did not have the RSN phenotype: PNA 98-11 (GenBank: QGTO01000003; C7426_103320-103334), PANS 02-1 (GenBank: NZ_QRDI01000007; C7423_RS14790-RS14860), and PANS 04-2 (GenBank: NZ_NMZV01000004; CG432_14985-15075) [RSN^-^/HiVir^+^] (Table 1). The resulting forward primer HiVir2pF (AATATCCATCAGTACCATT) overlapped with a T→C SNP conserved among the three previously described RSN^-^ strains within the *pepM* gene with the intention of reducing the rate of false positive PCR results (Figure 1A). The reverse primer HiVir2p_R (TGTTTAATGGGCCTTTTAC) annealed to a region in the *pavC* gene (GenBank: NZ_CP020943; B9Q16_20825).

**Table 1.**
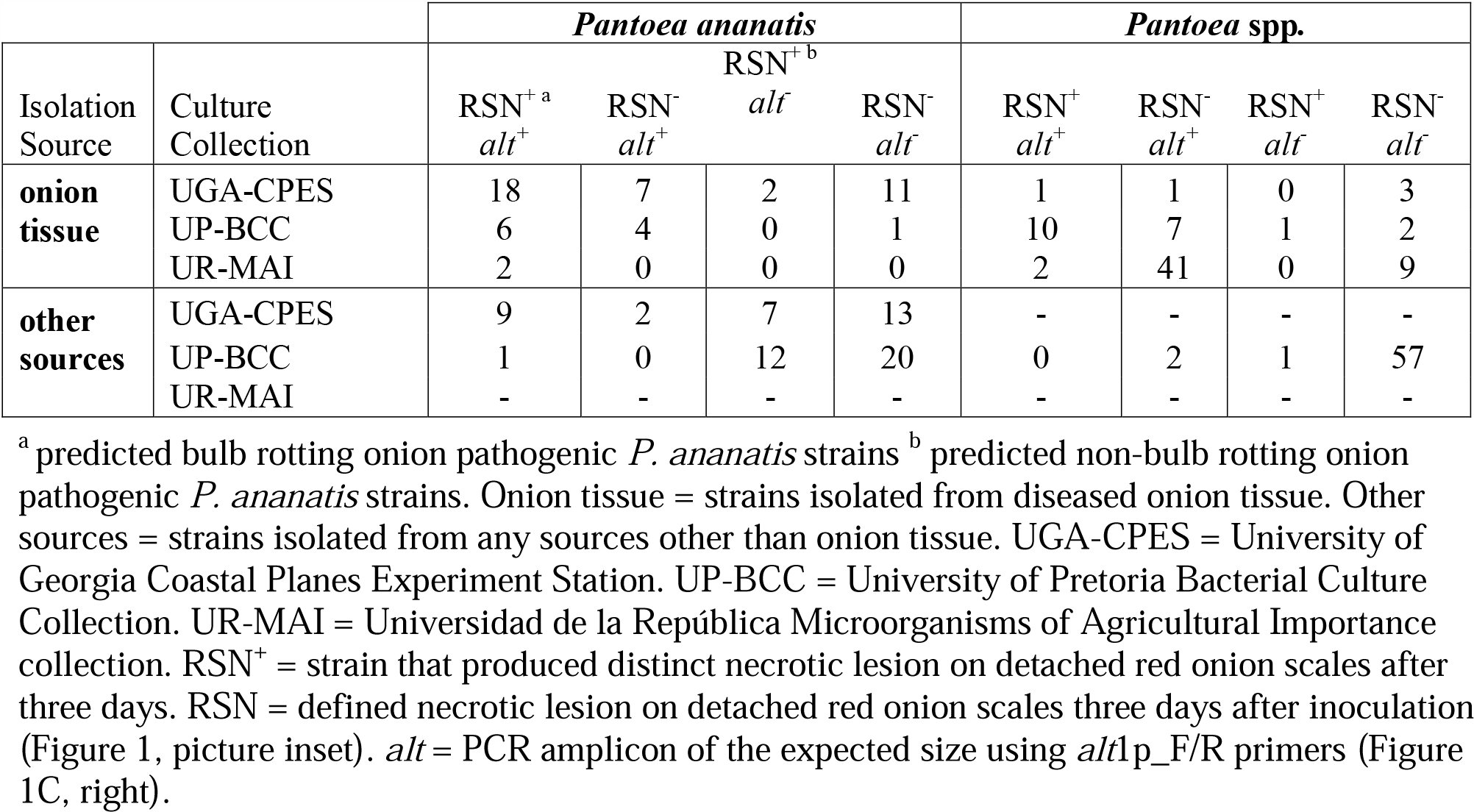
Summary of Red Scale Necrosis (RSN) and *alt* primer probe (*alt*1p F/R) results among bacterial culture collections (Supplementary Table 1).

**Figure 1.**
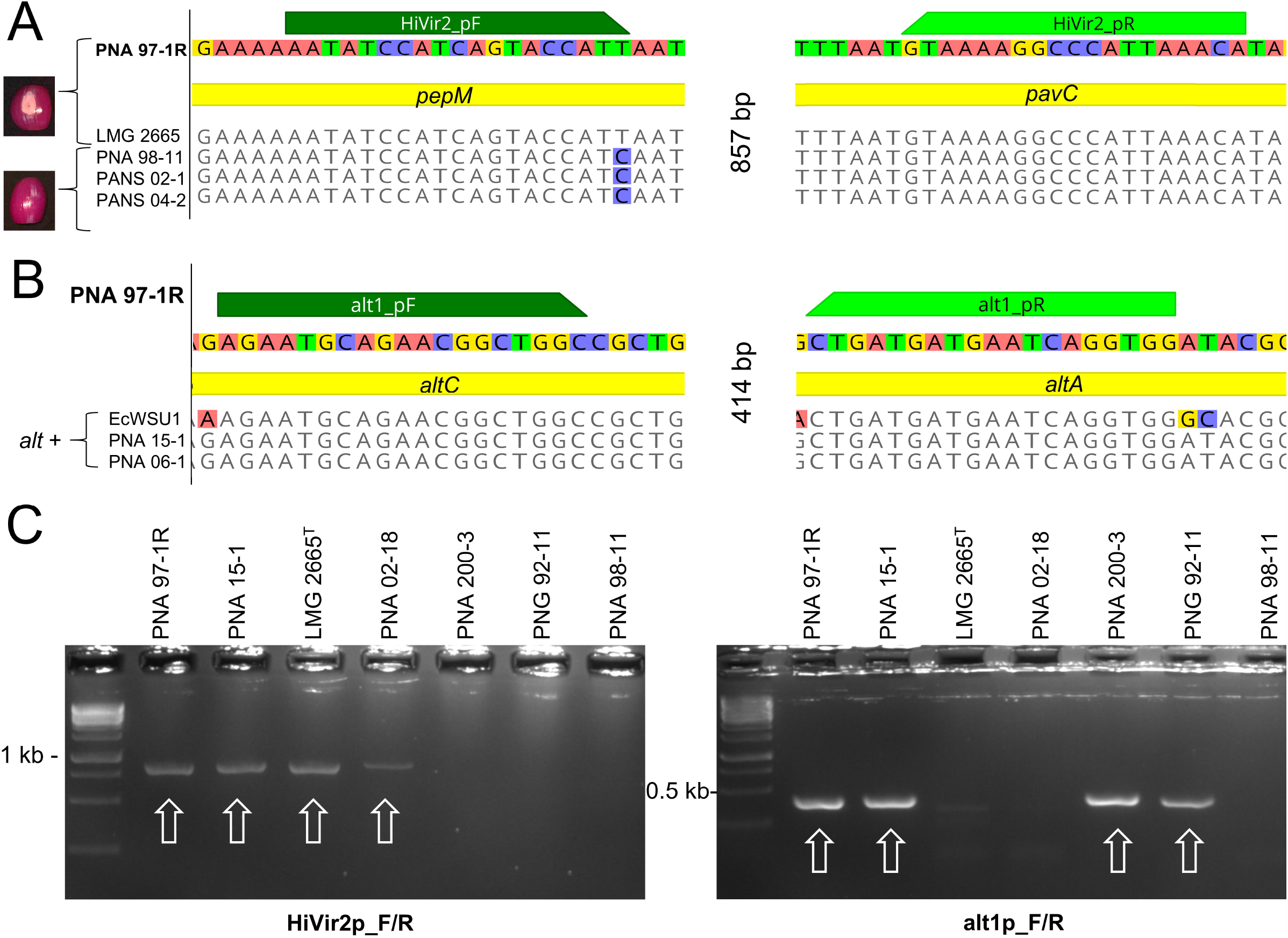
Design and testing of HiVir and *alt* primers. Design and testing of HiVir and *alt* primers. The HiVir cluster encodes proteins predicted for the production of a small molecule phosphonate phytotoxin. The *alt* cluster encodes proteins that encode tolerance to antimicrobial thiosulfinates produced in damaged onion tissues. (A) MAUVE alignment of sequenced *P. ananatis* strains with primer binding sites HiVir2_pF/R. All five strains possess the HiVir cluster; however, PNA 97-1R and LMG 2665^T^ induce red onion scale necrosis 3 days post inoculation while PNA 98-11, PNA 02-1, and PANS 04-2 do not (picture insets). HiVir2_pF primer incorporates the T→C SNP to aid in the detection of an RSN-associated HiVir cluster. (B) MAUVE alignment of sequenced *P. ananatis* and *E. ludwigii* strains with primer binding sites of *alt*1p_F/R. The three *P. ananatis* strains PNA 97-1R, PNA 15-1, and PNA 06-1 have functional *alt* clusters. *E. ludwigii* EcWSU1 contains a homologous gene cluster to generate inclusive primers. (C) Examples of gel electrophoresis of PCR amplicons. Strong amplicons of the predicted size were considered a positive PCR result. PCR assays included *P. ananatis* 97-1R as a HiVir and *alt* positive control. White arrows denote positive amplicons.

### 2.5 *alt* PCR assay design

The *alt*1p_F/R primer pair was designed to detect *P. ananatis* and *Enterobacter ludwigii* strains containing the *alt* gene cluster described by Stice *et al*. 2020. To identify conserved regions of the *alt* cluster a MAUVE alignment of three *P. ananatis* strains [PNA 15-1 (GenBank: NMZZ01000009; CG436_20695-20745), PNA 06-1 (GenBank: NMZY01000011; CG435_22460-22510), PNA 97-1R (GenBank: CP020945.2; B9Q16_23170-23120)] and one *E. ludwigii* strain [EcWSU1 (Genbank: CP002887.1; EcWSU1_A045-062)] was conducted (Figure 1B). The *alt*1p_F/R primers were generated from this alignment and the resulting primer pair, *alt*1p_F (AGAATGCAGAACGGCTGGC) and *alt*1p_R (CCACCTGATTCATCATCAG) were generated using the Primer3 (v2.3.7) plugin within Geneious Prime (v 2019.1.3).

### 2.6 PCR amplification

Primer assay oligos were ordered from Eurofins Genomics LLC (Louisville, KY). Primers were initially tested in a multiplex PCR reaction. However, to maintain consistency at multiple institutions, a simplex PCR scheme was adopted. Amplicon size and interpretation did not change when using a multiplex or simplex scheme. At UGA-CPES 74 strains were tested (0 multiplex, 74 simplex), at UP-BCC 54 strains were tested (42 multiplex, 82 simplex), and at UR-MAI 54 strains were tested (0 multiplex, 54 simplex). PCR reagents varied depending on the institution. Individual reactions conducted with UGA-CPES consisted of a 10 µL mixture containing 5 µL of GoTaq Green Master Mix 2X (Promega Corporation), 0.5 µL of template DNA, 0.5 µL of each primer and dH_2_O to the final volume. Reactions conducted with UP-BCC consisted of 10 μl reactions: 1 μl 10X DreamTaq Buffer, 0.5 μL of each primer, 0.4 μL 2 mM dNTPs, 0.5 μL genomic DNA, 0.2 µL DreamTaq DNA Polymerase (Thermo scientific), and dH_2_O to the final volume. Reactions conducted at UR-MAI consisted of 50 μl reactions: 5 μl 10X Standard Taq Buffer (NEB), 2.5 μL of each primer, 2 μL 10 mM dNTPs, 1 μL genomic DNA, and dH_2_O to the final volume. All reactions were run under the following conditions: 2 m at 98 °C; repeat 30 cycles of 10 s at 98 °C, 30 s at 47.4 °C (HiVir2p_F/R) / 52.3 °C (*alt*1p_F/R), 30 s at 72 °C; 10 m at 72 °C, hold at 4 °C. Amplified products were detected by gel electrophoresis of 10 μL of PCR reactions in a 1.5% TBE agarose gel with SYBR Safe dye (Thermo Scientific, Waltham, MA).

### 2.7 Determining the sensitivity of the *alt* and HiVir PCR assays

The sensitivity of each primer pair was tested using purified genomic DNA and a bacterial suspension of PNA 97-1R [HiVir^+^/*alt*^+^]. Purified genomic DNA (185 ng/µL) was serially diluted in 5-fold increments to 18,500; 3,700; 740; 148; 29.6; and 5.92 pg per PCR reaction and subjected to the simplex PCR assay described previously. A bacterial suspension of PNA 97-1R was prepared from a log-phase culture and adjusted to a concentration of 0.3 OD_600_ ≈ 1×10^8^ colony forming units (CFU)/mL. DNA was extracted using the boil-prep method previously described. The boiled 1×10^8^ CFU/mL suspension was diluted and added to individual reactions to obtain the following concentrations: 3,500; 3,000; 2,500; 2,000; 1,000; 500; 300; 200; 40; and 8 CFU per PCR reaction.

### 2.8 Validation of the HiVir and *alt* PCR assays using on whole genome sequencing data

Eighty-nine (74 UGA-CPES, 15 UP-BCC, 0 UR-MAI) of the strains tested with the *alt* and HiVir PCR assays have publicly available complete or draft whole genome sequences available (Supplementary Table 1). We determined the accuracy of the HiVir and *alt* PCR assays by comparing experimental PCR results to *in-silico* PCR results of sequenced genomes. Each individual genome was downloaded and imported into Geneious prime (v2019.1.3). The “test with saved primer” function was used to probe each primer pair against the genomes of the 89 sequenced strains. Results of individual virulence gene cluster presence by PCR assay in strains were tested for a significant association compared with the *in-silico* confirmation of *alt* and HiVir primer binding in sequenced genomes (*n*=89) by creating a 2×2 contingency table and calculating a two-tailed Fisher’s exact test in Microsoft Excel. The accuracy was calculated by summing up the true positive and true negative results and dividing by the total number of strains tested in this manner. A true positive is a strain that had a positive PCR amplicon and a positive *in-silico* primer bind. A true negative is a strain that had a negative PCR amplicon and a negative *in-silico* primer bind.

### 2.9 High-throughput red scale necrosis assay (RSN)

The red scale necrosis assay (RSN) was conducted as a high-throughput phenotypic assay to assess the onion pathogenic potential of strains in each culture collection. The assay was conducted as previously described with minor modifications (Stice et al., 2018). At UGA-CPES and UP-BCC, consumer produce red onions (*Allium cepa*. L.) were purchased, whereas at UR-MAI stored red onions cv. “Naqué” from postharvest experimental plots were taken, cut to approximately 3 cm wide scales, sterilized in a 3% household bleach solution for 1 m, promptly removed and rinsed in dH_2_O. Scales with a healthy unmarred appearance were used. Simple humidity chambers were used to encourage disease and prevent the drying of onion scales (approximately 80% relative humidity). Chambers consisted of a potting tray (27 × 52 cm) or similar plastic storage container, two layers of paper towels pre-wetted with distilled water, and the plastic removable portion of pipette trays or a similar item to prevent direct contact between the paper towels and the scales (Supplementary Figure 1). Disinfested onion scales were spaced evenly with approximately 1 cm of buffer between each scale in the humidity chamber (Supplementary Figure 1). Individual onion scales were wounded cleanly through the scale with a sterile pipette tip or needle and inoculated with a 10 μL drop of bacterial overnight LB or NA culture (Supplementary Figure 1). Sterile LB or NA culture was used as a negative control and strain PNA 97-1R was used as a positive control. The tray was covered with a plastic humidity dome or a loosely sealed container lid and incubated at room temperature for 72 h. Following incubation, strains that induced a distinct necrotic lesion with a defined border and clearing of the anthocyanin pigment on the onion scales were recorded as RSN^+^, while strains exhibiting no clearing were recorded as RSN^-^ (Figure 1A, picture inset).

### 2.10 Mini-Tn*7*Lux labeling of PNA 97-1R and LMG 2665^T^

To determine the colonization of the putative non-bulb rotting type strain *P. ananatis* LMG 2665^T^ [RSN^+^/HiVir^+^/*alt*^-^] and the known bulb rotting strain *P. ananatis* PNA 97-1R [RSN^+^/HiVir^+^/*alt*^+^], we used labeling methods as previously described (Stice et al., 2020). In brief, *Escherichia coli* (*Eco*) RHO3 pTNS3, *Eco* RHO5 pTn*7*PA143LuxFK, and the target *P. ananatis* strain were combined in a tri-parental mating. A five milliliter LB culture of each strain was grown O/N. After 12-14 h of incubation, 1 mL of each culture was centrifuged to pellet the bacteria. The supernatant was discarded, and the pellet resuspended in 100 µL of fresh LB broth. Twenty µL of each concentrated bacterial suspension was added to a sterile 1.5 mL microfuge tube. LB plates amended with 200 µg/mL diaminopimelic acid (DAP) were prepared, and sterile nitrocellulose membranes were placed on the plates. Amendment with DAP allows for growth of DAP-auxotrophic *Eco* RHO3 and RHO5 strains. A 20 µL volume of the mixed cultures and parental controls were spotted onto individual nitrocellulose membrane squares and allowed to dry. Following O/N incubation, the mixture was removed from the nitrocellulose membrane using a sterile loop and resuspended in 1 mL LB. The incubated mixed culture suspension was plated on Kanamycin (Km) LB selection plates. The following day Km-resistant colonies were selected and confirmed for luminescence with 2 m exposure settings with a ccd imager (analyticJena UVChemStudio, Upland, CA).

### 2.11 Neck-to-bulb infection assay

The neck-to-bulb infection assay was conducted as previously described (Stice et al., 2020). Three independent biological experiments were conducted with four technical repetitions per experiment. Three *alt* negative [RSN^+^/ HiVir^+^/*alt*^-^] strains [*P. ananatis* putative non-bulb rotting strains LMG 2665^T^ (causal agent of pineapple fruitlet rot, Tn7Lux), *P. ananatis* LMG 20103 (causal agent of Eucalyptus blight), and a Georgia *P. ananatis* natural variant lacking the *alt* gene cluster PNA 02-18 (onion origin)] were compared along with a negative dH_2_O control to three putative bulb-rotting alt-positive [RSN^+^/Hivir^+^/*alt*^+^] strains [PNA 97-1R (causal agent of onion center rot, Tn7Lux) and two additional Georgia strains isolated from onion in 2006 and 2015, PNA 15-1 and PNA 06-1 in a neck-to-bulb infection assay].

#### 2.11.1 Inoculum preparation

Cultures were grown O/N in LB, washed via centrifugation and resuspension in sterile dH_2_O, and the concentration adjusted to OD_600_ 0.3 ≈ 1×10^8^ CFU/mL in sterile dH_2_O with a final volume of 25 mL. Sterile toothpicks were soaked in the inoculum suspension for 20 m.

#### 2.11.2 Plant growth conditions and inoculation

At UGA-CPES, seven-week-old onion seedlings (cv. Sweet Agent) were obtained from the Vidalia Onion and Vegetable experiment station (Lyons, GA) and were potted individually in 16 cm × 15 cm (diameter × height) plastic pots with commercial potting mix on Dec 3^rd^, 2019. The onions were grown at the UGA South Milledge Greenhouse (Athens, GA). Greenhouse conditions were maintained at 24° C with 60% relative humidity and no supplemental lighting. Seedlings were watered using a hand hose directed at the base of the plants. Onion plants were inoculated at the end of the bulb development stage approximately 20 days before bulb maturation stage by inserting the inoculum-soaked toothpick horizontally through the onion neck just below the leaf fan (a favored thrips feeding site). Toothpicks were left in the plants. Colored tags were used to mark each treatment. Following inoculation, onions were randomized into blocks representing each replication. At 20 d post-inoculation, the onions were harvested for imaging. For harvesting, onion bulbs were removed from soil (which was sterilized and discarded following the experiment), rinsed with water, and cut twice transversely at the center of the bulb to produce a 1.5 cm section of the center of the onion (Supplementary Figure 2). The remaining portion of the top of the bulb had foliage removed and was cut longitudinally (Supplementary Figure 2).

#### 2.11.3 Imaging

Onions inoculated with non-labeled strains were imaged with a color camera and center rot symptom incidence was recorded. Onion inoculated with Tn*7*Lux auto-bioluminescent reporter strains (LMG 2665^T^ and PNA 97-1R) were imaged with a color camera followed by bright-field and long exposure imaging with the ccd imager (Analytik Jena UVP ChemStudio, Upland, CA). Within the VisionWorks software manual, long-exposure imaging was selected with the following settings: capture time 2 m, 70% focus, and 100% brightness (aperture), stack image, and saved to TIFF format. Brightfield images were captured with the following settings: capture time 40 mS, 70% focus, 60% brightness (aperture), and saved to TIFF format. After imaging, bright-field and long-exposure images were merged using ImageJ (Fiji release). The incidence of center rot symptoms and presence of a bioluminescent signal in long exposure images were recorded.

### 2.12 Statistical analysis

To determine whether a significant association exists between the HiVir PCR assay (positive vs. negative) and the RSN phenotype (positive vs. negative) we conducted a two-tailed Fisher exact test on 2×2 contingency tables in Microsoft Excel. The accuracy of a positive HiVir PCR assay as a predictor of the RSN phenotype was calculated by summing the true positive and true negative results and dividing by the total number of strains tested.

We tested whether a significant association exists between the *alt* PCR assay (positive vs. negative) and the source of isolation (onion vs. non-onion) using the two-tailed Fisher exact test previously described.

To determine whether source of isolation (onion or non-onion) or *Pantoea* group (*P. ananatis* or *Pantoea* spp.) was associated with an *alt*^*+*^ genotype and an RSN^+^ phenotype, we used a Z-proportions test to compare the proportion of four groups (RSN^+^ *alt*^-^, RSN^+^ *alt*^+^, RSN^-^ *alt*^+^, and RSN^-^ *alt*^-^). The two-tailed (α < 0.05) Z-proportions test assumes a normal distribution and considers the number of samples of each proportion being tested to determine whether the two proportions statistically differ according to a Z-statistic (Ramsey and Schafer, 2002).

To determine whether the percent center rot incidence at the onion midline differed among the three bulb rotting and non-bulb rotting strains, we conducted an ANOVA and Tukey-HSD test using R-Studio v1.2.1335 (package agricolae).

## 3 Results

An objective of this study was to assess the utility of two PCR assays and the phenotypic RSN assay for identifying *Pantoea* onion virulence genes. To accomplish this objective, we worked among three separate institutions to test the three assays. The assays were originally developed for use with *P. ananatis* strains; however, to investigate the breadth of their utility and potential limitations, we included a wider selection of *Pantoea* spp.

### 3.1 HiVir PCR assay

The HiVir cluster encodes predicted biosynthetic enzymes for the production of a small molecule phosphonate phytotoxin. The HiVir2p_F/R primer pair amplified an 857 bp portion of DNA including *pepM*, the intergenic region, and *pavC* while overlapping with an SNP associated with the RSN^-^ phenotype (Figure 1A, C). Amplification of the HiVir2p_F/R amplicon among all 252 strains tested in this study is depicted in Supplementary Table 1 (1=HiVir^+^, 0=HiVir^-^).

A positive HiVir PCR assay should be indicative of a present and functional HiVir cluster resulting in a positive RSN assay phenotype, while a negative HiVir PCR is expected to result in a negative RSN phenotype. The accuracy of a positive HiVir PCR assay corresponding to an *in-silico* primer bind was calculated among the 89 sequenced strains to be 95.51% with a significant association (*P*<0.001). The accuracy of PCR predicting an RSN^+^ phenotype was empirically determined among *P. ananatis* (*n*=115) and *Pantoea* spp. (*n*=137). The accuracy of the HiVir PCR assay among *P. ananatis* was 92.2 % and the accuracy among *Pantoea* spp. was 88.3 %. There was a significant association between the HiVir PCR result and the RSN result among *P. ananatis* (*P*<0.001) but not among *Pantoea*. spp. (*P*=0.296) (Supplementary Table 1).

### 3.2 *alt* PCR assay

The *alt* cluster encodes a cohort of putative disulfide exchange redox enzymes that confers tolerance to antimicrobial thiosulfinates produced in disrupted onion tissues. The *alt*1p_F/R primer pair amplified a 414 bp conserved portion of DNA encompassing the end of the *altB*, the intergenic region, and the beginning of *altC* (Figure 1B, C). Amplification of the *alt*1p_F/R amplicon is depicted in Supplementary Table 1 (1=*alt*^+^, 0=*alt*^-^).

The accuracy of a positive *alt* PCR assay corresponding to an *in-silico* primer bind was calculated among the 89 sequenced strains to be 95.51% with a significant association (*P*<0.001). There was a statistically significant association of the *alt* PCR assay (positive vs. negative) with isolation source (onion vs. non-onion) among *P. ananatis* (*n*=115; *P*<0.001) and *Pantoea* spp. (*n*=137; *P*<0.001) (Supplementary Table 1).

### 3.3 Sensitivity of PCR assays

Assessing the sensitivity of each PCR assay is important if these primers are used in the future for diagnostic purposes. The sensitivity of the assay refers to the minimum amount/concentration of DNA or bacterial template that can give a successful amplification. The minimum quantity of DNA with an interpretable amplicon was determined to be 29.6 pg of DNA per 10 µL reaction. The minimum number of colony forming units (CFU) per boil-prep template was determined to be 1,000 CFU for the HiVir2p_F/R primer pair and 300 CFU for the *alt*1p_F/R primer pair per 10 µL reaction.

### 3.4 Distribution of red scale necrosis (RSN) phenotype and HiVir *alt* PCR results

A positive HiVir amplicon was observed among 5 of the 17 species tested: *P. ananatis* (58/115), *P. allii* (1/21), *P. eucalypti* (2/39), *P. conspicua* (1/1), and *P. stewartii* subsp. *indologenes* (1/10) (Supplementary Table 1). Positive RSN results were observed among 6 of the 17 species within the *Pantoea* genus tested including: *P. ananatis* (55/115 tested), *P. agglomerans* (1/31 tested), *P. allii* (10/21 tested), *P. eucalypti* (4/39 tested), *P. stewartii* subsp. *indologenes* (1/10 tested), and *P. vagans* (1/5 tested) (Supplementary Table 1). A positive *alt* amplicon was observed in the same six species tested: *P. ananatis* (49/115 tested), *P. agglomerans* (13/31 tested), *P. allii* (18/21 tested), *P. eucalypti* (28/39 tested), *P. stewartii* subsp. *indologenes* (1/10 tested), and *P. vagans* (1/5 tested) (Supplementary Table 1).

### 3.5 Proportion of *P. ananatis* and *Pantoea* spp. strains among groups (onion vs. non-onion)

The proportion of *P. ananatis* strains from onion with RSN^+^ *alt*^+^ (26/51) was significantly greater than *P. ananatis-*types from non-onion sources (10/64) (*P<*0.001, Table 1, Figure 2, Supplementary Table 2). The proportion of *P. ananatis* from non-onion sources with RSN ^+^ *alt* ^-^ (19/64) was significantly greater compared to *P. ananatis* from onion (2/51) (*P*<0.001, Table 1, Figure 2, Supplementary Table 2). The proportion of *Pantoea* spp. strains from onion with RSN^-^ *alt*^+^ (49/77) was significantly greater compared to *Pantoea* spp. from non-onion sources (2/60) (*P*<0.001, Table 1, Figure 2, Supplementary Table 2).

**Figure 2.**
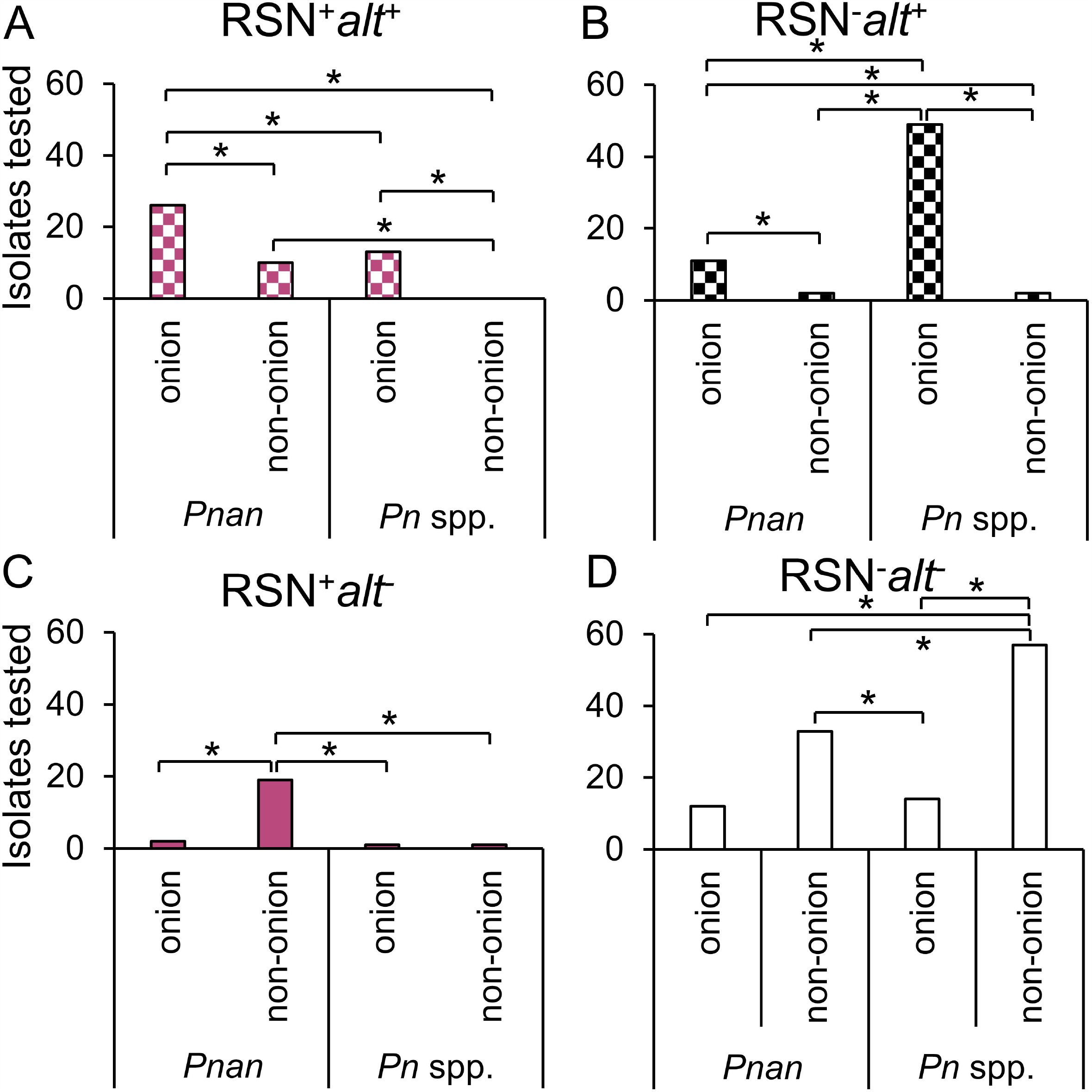
Distribution of RSN phenotype and *alt* PCR assay amongst *Pantoea* spp. Distribution of RSN phenotype and *alt* PCR assay amongst *Pantoea* spp. Each data point indicates a separate strain from Supplementary Table 1. RSN = defined necrotic lesion on detached red onion scales three days after inoculation (Figure 1, picture inset). *alt* = PCR amplicon of the expected size using *alt*1p_F/R primer (Figure 1C, right). Pnan = *Pantoea ananatis* strains. *Pn* spp. = all other *Pantoea* strains, excluding *P. ananatis*. Onion = stains isolated from diseased onion tissue. Non-onion = strains isolated from any sources other than onion tissue. A Z-statistic proportional analysis was performed among the defined groups. Brackets indicate a significant difference between two proportions. See Table 1 for raw counts. See Supplementary Table 2 for Z-statistic and P values.

### 3.6 *P. ananatis* neck-to-bulb infection assay

The percent incidence of dark necrotized tissue at the onion midline in the neck-to-bulb infection assay was significantly greater in *alt* positive [RSN^+^/Hivir^+^/*alt*^+^] *P. ananatis* strains PNA 97-1R (41.7%), PNA 15-1 (75%), and PNA 06-1(58.3%), compared to three *alt* negative [RSN^+^/Hivir^+^/*alt*^-^] *P. ananatis* strains LMG 2665^T^ (0%), LMG 20103 (0%), and PNA 02-18 (0%) (*P*<0.001, two-way ANOVA, HSD-post) (Figure 3A, B). Long exposure imaging of Tn*7*Lux autobioluminescent reporter strains LMG 2665^T^ and PNA 97-1R indicated a luminescence signal in the onion midline sections infected with PNA 97-1R that was absent in onions infected with LMG 2665^T^ (Figure 3A). This qualitative difference in onion bulb symptom production between the [RSN^+^/Hivir^+^/*alt*^-^] non-bulb-rotting *P. ananatis* type strain LMG 2665^T^ and [RSN^+^/Hivir^+^/*alt*^+^] bulb-rotting *P. ananatis* strain PNA 97-1R is consistent with previous evidence whereby the presence of the *alt* cluster in a HiVir^+^ strain leads to a higher incidence of bulb rot in the neck-to-bulb assay.

**Figure 3.**
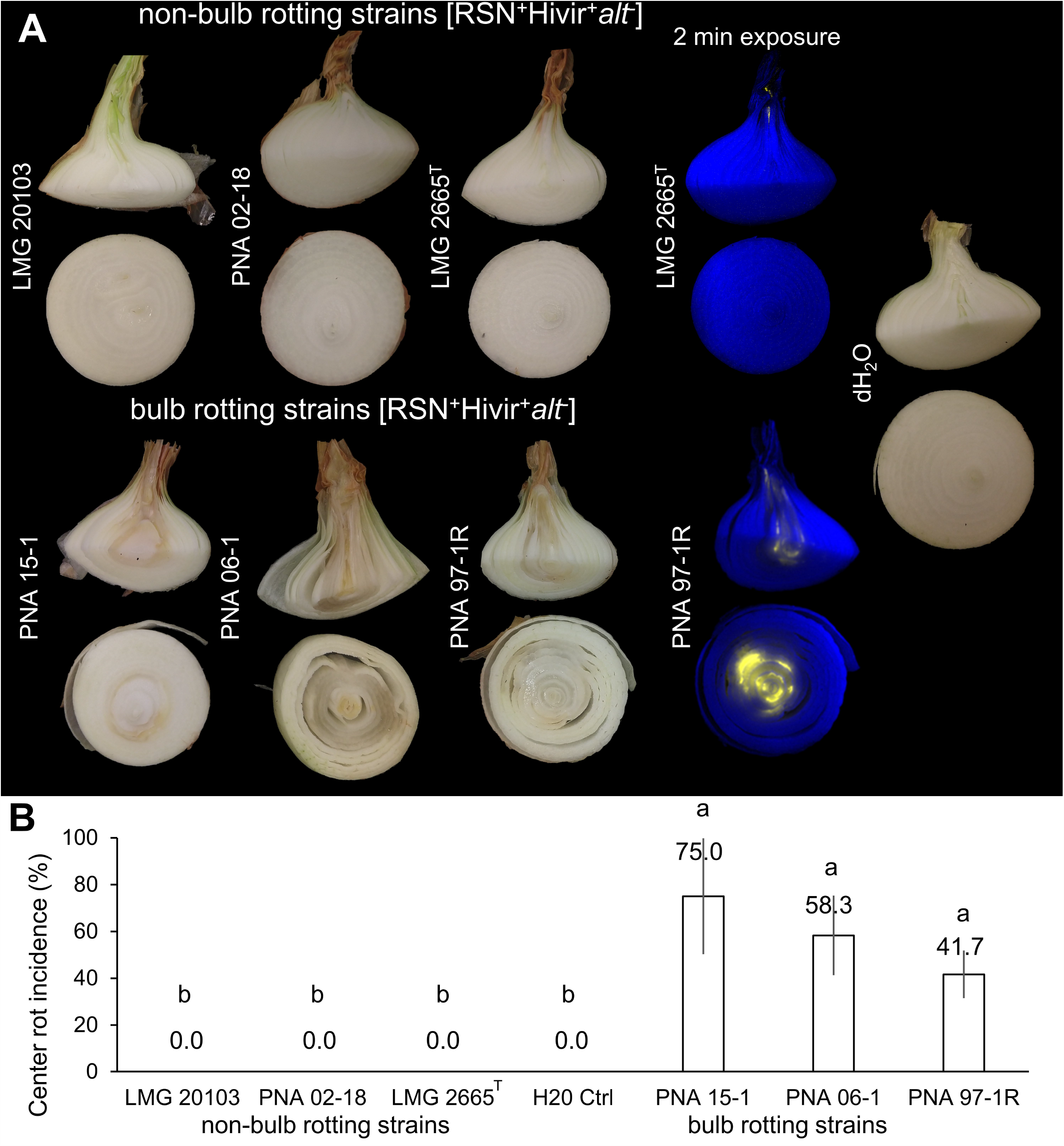
Neck-to-bulb infection assay correlates *P. ananatis* HiVir and *alt* traits with a bulb-rotting phenotype. Neck-to-bulb infection assay reveals bulb-rotting *P. ananatis* strains. Four onions were inoculated with a sterile toothpick soaked in a bacterial suspension of the respective strain 20 days prior to harvest in a randomized block design. Upon harvest, onions were cut at the onion midline, with the top portion cut again (Supplementary Figure 2). The incidence of a bioluminescent signal or tissue discoloration associated with center rot was recorded for the longitudinal and transverse sections. This experiment was conducted three times (n=12). (A) A panel of strains representing pathogenic but non-bulb-rotting *P. ananatis* strains [top] and pathogenic bulb-rotting *P. ananatis* strains [bottom]. The Tn*7*Lux labeled *P. ananatis* pathotype strain LMG 2665^T^ (causative agent of Eucalyptus blight) is compared to labeled PNA 97-1R (causative agent of center rot of onion), representative of the onion bulb rotting group and deposited to the international Belgium Coordinated Collection of Microorganisms under the accession LMG 31960.

## 4 Discussion

*Pantoea ananatis* is a broad host range pathogen with strains inducing disease symptoms ranging from fruitlet rots to leaf blights in a variety of hosts, including pineapple, eucalyptus, rice, maize, melon, and onion (Coutinho and Venter, 2009). To date, only two onion pathogen-specific virulence mechanisms have been genetically characterized: the chromosomal HiVir gene cluster and the plasmid localized *alt* gene cluster (Asselin et al., 2018; Stice et al., 2020). The HiVir gene cluster is hypothesized to code for biosynthesis of a yet unidentified phosphonate compound based on the genetic requirement for a characteristic *pepM* (phosphoenolpyruvate mutase) gene for *P. ananatis* to induce necrosis in onion tissue and tobacco leaves (Asselin et al., 2018). The *alt* gene cluster confers tolerance to the antimicrobial thiosulfinates released by damaged Allium tissues (Stice et al., 2020).

We developed two PCR assays to detect the HiVir and *alt* gene clusters and utilized the RSN assay to characterize the onion pathogenic potential of *P. ananatis* strains. We applied these assays to determine the distribution of onion-associated virulence gene clusters among *Pantoea* strains in three unique culture collections comprised of 252 strains. Previous studies have identified that the HiVir and *alt* gene clusters are distributed outside of *P. ananatis*, as demonstrated by their presence in strains of *P. stewartii* subsp. *indologenes, P. allii*, and *P. agglomerans* based on whole genome sequencing (Stice et al., 2020). By leveraging these simple assays against the UGA-CPES (focused on diseased onion strains, primarily of *P. ananatis*, or isolated from the onion-associated weeds and thrips) and UP-BCC (focused on many *Pantoea* spp. including pathogenic rice, maize, and eucalyptus strains, among others), we observed a significant enrichment of the *alt* gene cluster among strains recovered from diseased onions, and a strong statistical association between the HiVir PCR and RSN phenotypic assays (Figure 2, Supplementary Table 1). The distribution of these onion virulence genetic clusters will be useful for future studies of *Pantoea* strains involved in the onion center rot complex. In addition, defining genetic features of bulb-rotting *P. ananatis* strains, based on the presence or absence of the *alt* cluster, will help distinguish strains of *P. ananatis* that are a greater threat to onion production (Figure 3).

### 4.1 Utility of HiVir PCR assay

We designed the HiVir PCR assay to detect *P. ananatis* strains that induce red onion scale necrosis. To do this, we compared three strains in our collection that have the HiVir cluster in their sequenced draft genomes but lack the RSN phenotype. The forward primer, HiVir2p_F, overlaps with a conserved SNP among the RSN^-^/HiVir^+^ strains. Our results indicate the HiVir2p_F/R primers work as expected with no amplification in the RSN^+^/HiVir^-^ strains such as PNA 98-11 (Figure 1A,C). The HiVir PCR assay is sensitive and accurate in predicting the RSN ability of *P. ananatis* strains with a significant association between the PCR result and the RSN phenotype. Among *Pantoea* spp. (excluding *P. ananatis*) the HiVir PCR assay is less accurate and does not have a statistically significant association with RSN. An example would be *P. vagans* LMG 24196, a strain isolated from eucalyptus leaves and shoots showing symptoms of blight and dieback in Argentina (Brady et al., 2009). LMG 24196 is RSN^+^ but HiVir^-^ in our tests. We hypothesize that LMG 24196 either possesses a divergent, non-*P. ananatis*-type HiVir cluster that is not amplified by our HiVir primer set, or that this strain may use an alternative virulence mechanism to cause red scale necrosis. Seventeen of the 252 strains sampled in this study display an RSN^+^ HiVir^-^ result (1 UGA-CPES, 14 UP-BCC, 2 UR-MAI). Due to the weak correlation between the HiVir PCR assay and the RSN phenotype outside of *P. ananatis*, we have concluded that RSN phenotyping is the more reliable assay for determining the onion pathogenic potential of strains both within *P. ananatis* and among *Pantoea* spp. With this information, we organized and grouped our results from Supplementary Table 1 to focus on the RSN phenotype for most figures and tables in this study.

### 4.2 Utility of RSN assay

The RSN assay may be predictive of the general plant pathogenic potential of *Pantoea* strains infecting hosts other than onion such as maize and rice. *P. ananatis* BD 647 was isolated from symptomatic maize, and its pathogenicity was experimentally validated; in this study it tested RSN^+^/HiVir^+^/*alt*^-^ (Goszczynska et al., 2007). *P. agglomerans* LMG 2596 was isolated from a symptomatic onion umbel, and its pathogenicity was validated; in this study it tested RSN^+^/HiVir^-^ /*alt*^*+*^ (Hattingh and Walters, 1981). *P. ananatis* strains causing stem necrosis and palea browning of rice, including DAR76141, CTB1135, SUPP2113, had an RSN^+^/HiVir^-^/*alt*^*-*^ result (Table S1) (Cother et al., 2004; Kido et al., 2008, 2010).

### 4.3 Utility of *alt* PCR assay

We designed the *alt* PCR assay to detect the *alt* cluster among *P. ananatis* strains and a homologous cluster present in an *Enterobacter ludwigii* strain as described in our previous work (Stice et al., 2020). The *alt* cluster confers tolerance to antimicrobial thiosulfinates that are present in onion and garlic tissues. Due to the reactive and unstable nature of these molecules, it was not practical to test all strains for thiosulfinate tolerance in a high throughput manner. However, we tested for and found a statistically significant association between a positive *alt* PCR assay and strains sourced from onion tissue in both *P. ananatis* strains and *Pantoea* spp. (Table 1, Figure 2).

### 4.4 Distribution of onion associated virulence gene clusters

#### 4.4.1 UGA-CPES

The UGA-CPES collection included mostly *P. ananatis* strains isolated from symptomatic onions along with strains isolated from onion-associated weeds and thrips. *P. ananatis* strains originating from symptomatic onion typically contained the RSN^+^/HiVir^+^/*alt*^*+*^ result; however, strains with only HiVir, only *alt*, or neither gene cluster were also recovered. We would expect center-rot-causing strains to harbor both traits, as these virulence determinants have been demonstrated experimentally to strongly contribute to neck-to-bulb infection. Among *P. ananatis* strains isolated from sources other than onion, the RSN^+^/*alt*^*-*^ result was more pronounced (Table 1).

#### 4.4.2 UP-BCC

The UP-BCC collection included *Pantoea* strains from diverse sources with specific enrichment in maize, eucalyptus, and onion. Most strains tested RSN^-^*alt*^*-*^ (Table 1). *P. ananatis* species had enrichment of the RSN^+^ and *alt*^*+*^ results when isolated from onion. There was a larger proportion of *P. ananatis* strains in the UP-BCC with the RSN^+^*alt*^*-*^ phenotype, which supports our interpretation that RSN^+^/HiVir^+^ may be broadly associated with *P. ananatis* plant pathogenicity but that the *alt* cluster is common among strains associated with onion.

#### 4.4.3 UR-MAI

The UR-MAI collection was entirely comprised of strains originating from diseased onion. The RSN^+^/HiVir^+^/*alt*^*+*^ result was obtained from the two strains belonging to the species *P. ananatis* within this collection. Interestingly, the RSN^+^/*alt*^+^ result was also found in one *P. eucalypti* strain (MAI 6036) and one *P. agglomerans* strain (MAI6045) recovered from symptomatic onion leaves and seed stalks. The HiVir^-^/RSN^-^/*alt*^+^ result was nearly universally distributed among recovered strains from this survey (Supplementary Table 1, Figure 2, Table 1). This suggests that alternative onion necrosis mechanisms exist among *Pantoea* spp.

### 4.5 Proportional analysis of RSN and *alt* results among strains

Considering the selection of strains was biased in regard to isolation source (mostly of onion origin), we grouped the RSN/*alt* results of our strains based on two metrics for a proportional analysis: 1.) onion vs. non-onion strains, 2.) *P. ananatis* vs. *Pantoea* spp. strains. We then conducted a *Z*-proportions test. This test allowed us to reveal broad differences by comparing proportions of strains in defined groups. A significantly greater portion of strains of onion origin have the RSN^+^/*alt*^*+*^ and RSN^-^/*alt*^+^ results compared to those of non-onion origin for both *P. ananatis* and *Pantoea* spp. strains (Figure 2, Supplementary Table 2). This offers support for our conceptual model of *alt* cluster enrichment among onion pathogenic strains. Conversely, a significantly greater proportion of *P. ananatis* strains of non-onion origin has the RSN^+^/*alt*^*-*^ result compared to those of onion origin (Figure 2, Supplementary Table 2). This suggests that strains causing foliar symptoms on crops such as rice and maize may utilize the HiVir encoded toxin but do not require the *alt* cluster, as their hosts do not produce the thiosulfinate defensive chemistry found in onions.

### 4.6 The association of HiVir and *alt* gene clusters with *P. ananatis* bulb rot symptoms

We previously demonstrated that both the HiVir and *alt* gene clusters were required by *P. ananatis* PNA 97-1R [RSN^+^/HiVir^+^/*alt*^+^] to create center rot associated symptoms in a neck-to-bulb infection assay (Stice et al., 2020). We expected the *alt* positive putative bulb-rotting strains PNA 97-1R, PNA 15-1, and PNA 06-1 [RSN^+^/HiVir^+^/*alt*^+^] to cause symptomatic bulb rots at the onion midline compared to *alt* negative putative non-bulb-rotting strains LMG 2665^T^ (causal agent of pineapple fruitlet rot), PNA 02-18 (natural onion isolate lacking *alt*), and LMG 20103 (causal agent of eucalyptus blight) [RSN^+^/HiVir^+^/*alt*^-^]. Our results were consistent with this hypothesis whereby the three natural *alt* positive [RSN^+^/Hivir^+^/*alt*^+^] *P. ananatis* strains had a significantly higher incidence of bulb rot symptoms in the neck-to-bulb infection assay compared to *alt* negative [RSN^+^/Hivir^+^/*alt*^-^] *P. ananatis* strains (Figure 3). To facilitate access to the *alt* positive [RSN^+^/Hivir^+^/*alt*^-+^] *P. ananatis* strain PNA 97-1R, we have deposited it to the international Belgium Coordinated Collection of Microorganisms under the accession LMG 31960.

## 4.7 Conclusion

The three culture collections tested in this study have allowed us to gather a preliminary picture of the distribution of the [HiVir^+^ /*alt*^+^] genotypes and RSN^+^ phenotype associated with onion center rot. The HiVir and *alt* PCR assays which were respectively designed to be specific and inclusive were validated on a panel of sequenced genomes with 95.15% accuracy (Figure 1). The RSN assay was confirmed to be a robust method capable of assessing *Pantoea* strains carrying pathogenic potential (Figure 2). A positive RSN and HiVir PCR assay were significantly associated among *P. ananatis* strains; however, there was not a significant association among *Pantoea* spp., suggesting the assay could be improved by including new sequencing information in future primer designs. Conversely, the *alt* PCR assay allowed for the detection of *alt* associated genes in six *Pantoea* species with a significant correlation between a positive *alt* PCR assay and strains of onion tissue origin among both *P. ananatis* and *Pantoea* spp. We also demonstrated that natural *alt* positive [RSN^+^/HiVir^+^/*alt*^*+*^] *P. ananatis* congeners have enhanced bulb rot potential in the neck-to-bulb infection assay compared to *alt* negative [RSN^+^/HiVir^+^/*alt*^*-*^] *P. ananatis* congeners.

## 5 Conflict of Interest

The authors declare that the research was conducted in the absence of any commercial or financial relationships that could be construed as a potential conflict of interest.

## 6 Author Contributions

S.S., B.K., and B.D. conceived and planned the experiments in this study. S.S. designed and tested primers. S.S. and S.K. tested the UGA-CPES strain collection. S.S., G.S., and T.C. tested the UP-BCC strain collection. S.A., G.G., and M.S. tested the UR-MAI strain collection. S.S. completed auto-bioluminescent strain labeling and the bulb-to-scale pathogenicity assay. S.S. processed experimental data and designed figures. P.S. advised and completed statistical analyses. All authors discussed and contributed to the final manuscript.

## 7 Funding

This work was supported with funding from the Vidalia Onion Committee to B.K. and B.D., United States Department of Agriculture (USDA) SCBGP project AM180100XXXXG014 to B.D. We acknowledge support from grants FMV 104703 (ANII, Uruguay) and CSIC Grupos de Investigación I+D 2000 (CSIC, Udelar, Uruguay) to S.D.A., G.A.G. and M.I.S. This work is supported by Specialty Crops Research Initiative Award 2019-51181-30013 from the USDA National Institute of Food and Agriculture to B.D. and B.K. Any opinions, findings, conclusions, or recommendations expressed in this publication are those of the author(s) and do not necessarily reflect the view of the U.S. Department of Agriculture.

## 8 Acknowledgments

We would like to acknowledge Li Yang for use of equipment, and members of the Yang and Kvitko Labs, as well as Ron Walcott and Brenda Schroeder, for helpful discussions regarding the preparation of the manuscript.

## Notes

### Competing Interest Statement

The authors have declared no competing interest.

