## Supplementary Table 1, Supplementary Table 2, Supplementary Figure 2, Supplementary Figure 1 for "The distribution of onion virulence gene clusters among *Pantoea* spp"

Supplementary Material

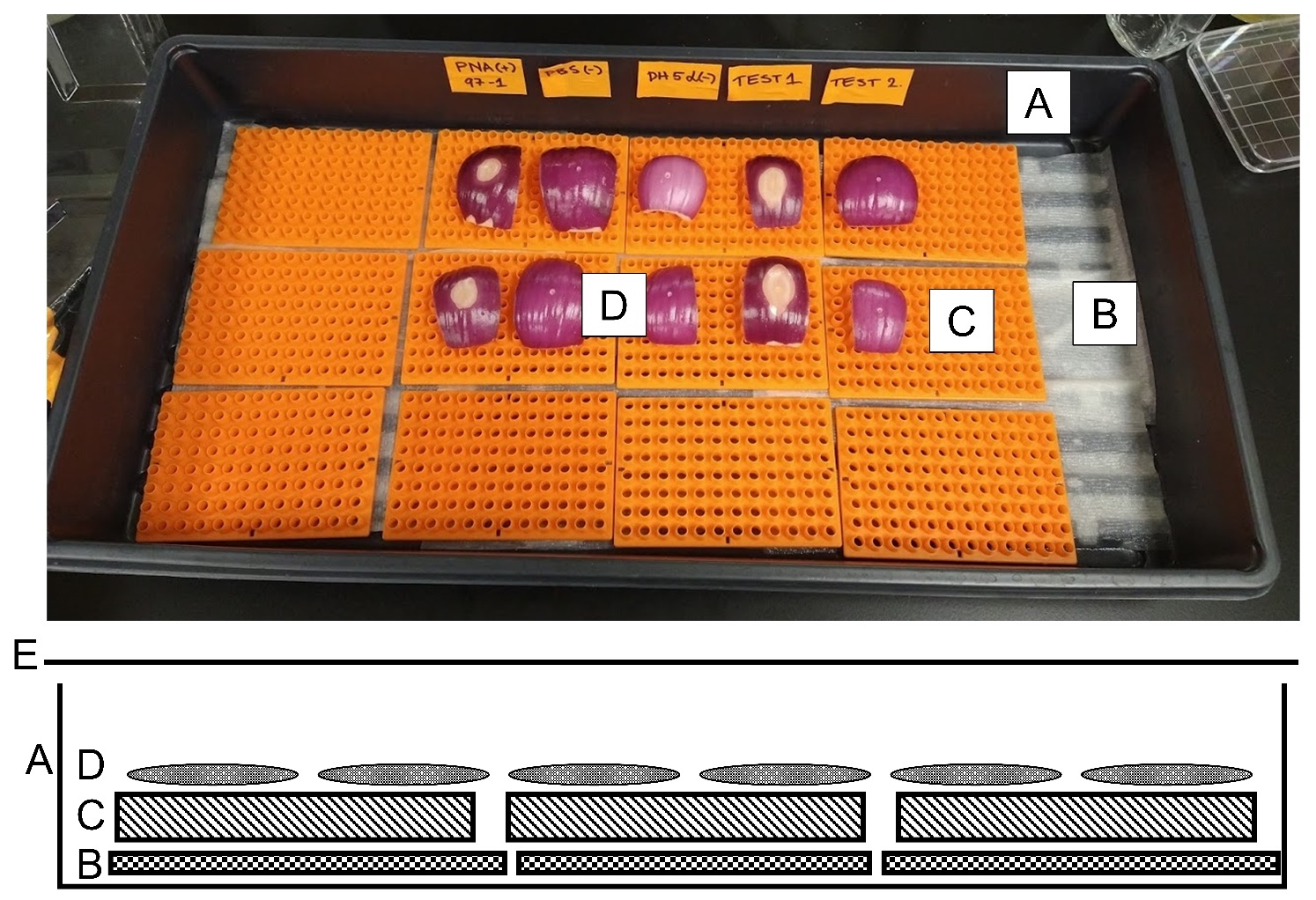

**Supplementary Figure 1.** High-throughput red scale necrosis assay. (Top) humid-chamber set-up for UGA-CPES assay. (Bottom) Graphical representation of humid-chamber set-up. (A) Plastic potting tray (27 × 52 cm) or plastic container (B) Pre-moistened paper towels (C) Plastic removable portion of 20 μl pipette trays or similar to prevent direct contact between scales and paper towels (D) Bleach disinfested red onion scales (E) Plastic humidity dome or loose fitting plastic cover. Detached red onion scales are disinfested, wounded, inoculated with a 10 µL bacterial suspension, and assessed for the red scale necrosis phenotype (PNA 97-1, Test1) after 3 days.

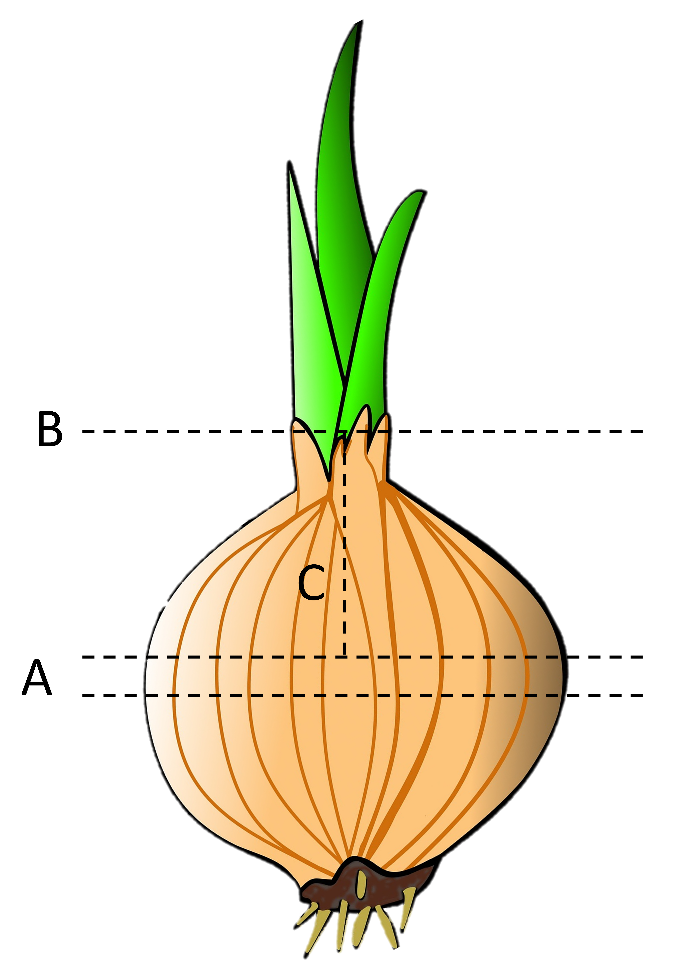

**Supplementary Figure 2.** Dissection of onions in systemic onion pathogenicity assay. (A) Onions were cut twice transversely at the center of the onion to produce a 1.5 cm section of the center of the onion. (B) Foliage removed. (C) Remaining top portion of bulb cut longitudinally. See figure 3 for images of cut onions.

**Supplementary Table 1.** Strains tested for Red Scale Necrosis and virulence gene clusters (*alt* / HiVir primer assay).

| **Species** | **Strain** | **Source** | **Location [Country (State/Region) city/subregion]** | **Year** | **GenBank Accession** | **Publication** | **Red Scale Necrosis** | **HiVir2p F/R** | **alt1p F/R** |
| --- | --- | --- | --- | --- | --- | --- | --- | --- | --- |
| *P. ananatis* ^A^ | PNA 97-1R ^I^ | onion | U.S.A. (Georgia) Toombs | 1997 | CP020943.2 - CP020945.2 | Gitaitis & Gay 1997 | 1 | 1 | 1 |
| *P. ananatis* ^A^ | PNA 99-7 ^I^ | onion | U.S.A. (Georgia) Tattnall | 1999 | NMZW00000000 | Stice et al. 2018 | 0 | 0 | 0 |
| *P. ananatis* ^A^ | PANS 99-23 ^I^ | yellow nutsedge | U.S.A. (Georgia) Terrell | 1999 | NMZS00000000 | Stice et al. 2018 | 0 | 0 | 0 |
| *P. ananatis* ^A^ | PANS 99-36 ^I^ | florida pusley | U.S.A. (Georgia) Tifton | 1999 | NMZT00000000 | Stice et al. 2018 | 0 | 0 | 0 |
| *P. ananatis* ^A^ | PANS 04-2 ^I^ | tobacco thrips | U.S.A. (Georgia) Toombs | 2004 | NMZV00000000 | Stice et al. 2018 | 0 | 0 | 0 |
| *P. ananatis* ^A^ | PNA 15-1 ^I^ | onion | U.S.A. (Georgia) Tattnall | 2015 | NMZZ00000000 | Stice et al. 2018 | 1 | 1 | 1 |
| *P. ananatis* ^A^ | PNA 200-3 ^I^ | onion | U.S.A. (Georgia) Tifton | 2000 | NMZX00000000 | Stice et al. 2018 | 0 | 0 | 1 |
| *P. ananatis* ^A^ | PANS 99-3 ^I^ | florida pusley | U.S.A. (Georgia) Tifton | 1999 | NMZR00000000 | Stice et al. 2018 | 1 | 1 | 1 |
| *P. ananatis* ^A^ | PANS 01-2 ^I^ | onion thrips | U.S.A. (Georgia) Tifton | 2001 | NMZU00000000 | Stice et al. 2018 | 1 | 1 | 1 |
| *P. ananatis* ^A^ | PNA 06-1 ^I^ | onion | U.S.A. (Georgia) Wayne | 2006 | NMZY00000000 | Stice et al. 2018 | 1 | 1 | 1 |
| *P. ananatis* ^A^ | PNA 200-7 ^I^ | onion | U.S.A. (Georgia) Tifton | 2000 | QGGN00000000 | this paper | 1 | 1 | 1 |
| *P. ananatis* ^A^ | PNA 07-10 ^I^ | onion | U.S.A. (Georgia) Toombs | 2007 | QTTO00000000 | this paper | 1 | 1 | 1 |
| *P. ananatis* ^A^ | PNA 02-18 ^I^ | onion | U.S.A. (Georgia) Tattnall | 2002 | RBXY00000000 | Stice et al. 2020 | 1 | 1 | 0 |
| *P. ananatis* ^A^ | PANS 200-1 ^I^ | slender amaranth | U.S.A. (Georgia) Lyons | 2000 | QTTV00000000 | this paper | 0 | 0 | 0 |
| *P. ananatis* ^A^ | PNA 07-1 ^I^ | onion | U.S.A. (Georgia) Tifton | 2007 | QICU00000000 | this paper | 1 | 1 | 1 |
| *P. ananatis* ^A^ | PANS 02-01 ^I^ | tobacco thrips | U.S.A. (Georgia) Tattnall | 2002 | QRDI01000000 | this paper | 0 | 0 | 0 |
| *P. ananatis* ^A^ | PNA 86-1 ^I^ | peach | U.S.A. (Georgia) - | 1986 | QLSY00000000 | this paper | 0 | 0 | 0 |
| *P. ananatis* ^A^ | PNA 98-11 ^I^ | onion | U.S.A. (Georgia) Evans | 1998 | QGTO00000000 | this paper | 0 | 0 | 0 |
| *P. ananatis* ^A^ | PNA 14-1 ^I^ | onion | U.S.A. (Georgia) - | 2014 | QEKS00000000 | this paper | 0 | 0 | 1 |
| *P. ananatis* ^A^ | PNA 11-1 ^I^ | onion | U.S.A. (Georgia) Lyons | 2011 | QGTK00000000 | this paper | 0 | 0 | 0 |
| *P. allii* ^A^ | PNA 200-10 ^I^ | onion | U.S.A. (Georgia) - | 2000 | QGHF00000000 | this paper | 0 | 0 | 0 |
| *P. agglomerans* ^A^ | PNG 97-1 ^I^ | onion | U.S.A. (Georgia) - | 1997 | SOSD00000000 | this paper | 0 | 0 | 0 |
| *P. agglomerans* ^A^ | PNG 92-11 ^I^ | onion | U.S.A. (Georgia) - | 1992 | QGHE00000000 | this paper | 0 | 0 | 1 |
| *P. stewartii* subsp. *indologens* ^A^ | PNA 14-12 ^I^ | onion | U.S.A. (Georgia) Vidalia | 2014 | SOAJ00000000 | Stumpf et al. 2017 | 0 | 0 | 0 |
| *P. stewartii* subsp*. indologens* ^A^ | PNA 03-3 ^I^ | onion | U.S.A. (Georgia) Toombs | 2003 | QICO00000000 | Stumpf et al. 2017 | 1 | 1 | 1 |
| *P. ananatis* ^B^ | PANS 1-5 ^I^ | tobacco thrips | U.S.A (Georgia) Tifton | 2001 | - | this paper | 1 | 1 | 1 |
| *P. ananatis* ^B^ | PNA 18-5S ^I^ | onion | U.S.A (Georgia) Vidalia | 2018 | - | this paper | 1 | 1 | 1 |
| *P. ananatis* ^B^ | PANS 1-6 ^I^ | thrips | U.S.A (Georgia) Tifton | 2001 | - | this paper | 1 | 1 | 1 |
| *P. ananatis* ^B^ | PANS 2-5 ^I^ | thrips from peanut | U.S.A (Georgia) Tifton | 2002 | - | this paper | 1 | 1 | 0 |
| *P. ananatis* ^B^ | PNA 18-7S ^I^ | onion | U.S.A (Georgia) Tifton | 2018 | - | this paper | 1 | 1 | 1 |
| *P. ananatis* ^B^ | PNA 18-1 ^I^ | onion | U.S.A (Georgia) Tifton | 2018 | - | this paper | 1 | 1 | 1 |
| *P. ananatis* ^B^ | PNA 6-1 ^I^ | onion | U.S.A (Georgia) Wayne | 2006 | - | this paper | 1 | 1 | 1 |
| *P. ananatis* ^B^ | PNA 99-2 ^I^ | onion | U.S.A (Georgia) Tattnall | 1999 | - | this paper | 1 | 1 | 1 |
| *P. ananatis* ^B^ | PNA 7-10 ^I^ | onion | U.S.A (Georgia) Toombs | 2007 | - | this paper | 1 | 1 | 1 |
| *P. ananatis* ^B^ | PNA 18-9S ^I^ | onion | U.S.A (Georgia) Vidalia | 2018 | - | this paper | 1 | 0 | 1 |
| *P. ananatis* ^B^ | PANS 2-6 ^I^ | thrips from peanut | U.S.A (Georgia) Tifton | 2002 | - | this paper | 1 | 1 | 1 |
| *P. ananatis* ^B^ | PNA 7-1 ^I^ | onion | U.S.A (Georgia) Tattnall | 2007 | - | this paper | 1 | 1 | 1 |
| *P. ananatis* ^B^ | PANS 99-33 ^I^ | florida pusley | U.S.A (Georgia) Coffee | 1999 | - | this paper | 1 | 1 | 0 |
| *P. ananatis* ^B^ | PANS 99-25 ^I^ | bristly starbur | U.S.A (Georgia) Vidalia | 1999 | - | this paper | 1 | 1 | 0 |
| *P. ananatis* ^B^ | PNA 18-2 ^I^ | onion | U.S.A (Georgia) Tifton | 2018 | - | this paper | 1 | 1 | 1 |
| *P. ananatis* ^B^ | PNA 03-2 ^I^ | onion | U.S.A (Georgia) Tifton | 2003 | - | this paper | 0 | 0 | 1 |
| *P. ananatis* ^B^ | PANS 99-32 ^I^ | florida pusley | U.S.A (Georgia) Vidalia | 1999 | - | this paper | 0 | 1 | 0 |
| *P. ananatis* ^B^ | PNA 98-3 ^I^ | onion | U.S.A (Georgia) Dougherty | 1998 | - | this paper | 0 | 0 | 0 |
| *P. ananatis* ^B^ | PNA 13-1 ^I^ | onion | U.S.A (Georgia) Lyons | 2013 | - | this paper | 0 | 0 | 0 |
| *P. ananatis* ^B^ | PNA 14-2 ^I^ | onion | - | 2014 | - | this paper | 0 | 0 | 0 |
| *P. ananatis* ^B^ | PNA 15-3 ^I^ | onion | U.S.A (Georgia) Tattnall | 2015 | - | this paper | 0 | 0 | 0 |
| *P. ananatis* ^B^ | PNA 18-10 ^I^ | onion | U.S.A (Georgia) Vidalia | 2018 | - | this paper | 0 | 0 | 1 |
| *P. ananatis* ^B^ | PNA 18-6S ^I^ | onion | U.S.A (Georgia) Vidalia | 2018 | - | this paper | 0 | 0 | 1 |
| *P. ananatis* ^B^ | PNA 18-8S ^I^ | onion | U.S.A (Georgia) Vidalia | 2018 | - | this paper | 0 | 0 | 1 |
| *P. ananatis* ^B^ | PNA 200-1 ^I^ | onion | U.S.A (Georgia) Toombs | 2000 | - | this paper | 0 | 0 | 1 |
| *P. ananatis* ^B^ | PNA 200-8 ^I^ | onion | U.S.A (Georgia) Tifton | 2000 | - | this paper | 0 | 0 | 1 |
| *P. ananatis* ^B^ | PANS 99-5 ^I^ | prairie verbena | U.S.A (Georgia) Tifton | 1999 | - | this paper | 0 | 0 | 0 |
| *P. ananatis* ^B^ | PANS 99-4 ^I^ | florida pusley | U.S.A (Georgia) Tifton | 1999 | - | this paper | 1 | 1 | 1 |
| *P. ananatis* ^B^ | PANS 200-2 ^I^ | pink purslane | U.S.A (Georgia) Tifton | 2000 | - | this paper | 1 | 1 | 1 |
| *P. ananatis* ^B^ | PANS 99-29 ^I^ | crab grass | U.S.A (Georgia) Tifton | 1999 | - | this paper | 1 | 1 | 0 |
| *P. ananatis* ^B^ | PANS 2-7 ^I^ | thrips from peanut | U.S.A (Georgia) Tifton | 2002 | - | this paper | 1 | 1 | 0 |
| *P. ananatis* ^B^ | PNA 18-5 ^I^ | onion | U.S.A (Georgia) Vidalia | 2018 | - | this paper | 1 | 1 | 1 |
| *P. ananatis* ^B^ | PNA 99-14 ^I^ | onion | U.S.A (Georgia) Tattnall | 1999 | - | this paper | 1 | 1 | 1 |
| *P. ananatis* ^B^ | PANS 1-2 ^I^ | thrips from onion | U.S.A (Georgia) Tifton | 2001 | - | this paper | 1 | 1 | 1 |
| *P. ananatis* ^B^ | PNA 99-9 ^I^ | onion | U.S.A (Georgia) Tattnall | 1999 | - | this paper | 1 | 1 | 1 |
| *P. ananatis* ^B^ | PNA 18-3S ^I^ | onion | U.S.A (Georgia) Vidalia | 2018 | - | this paper | 1 | 1 | 0 |
| *P. ananatis* ^B^ | PANS 2-8 ^I^ | thrips from peanut | U.S.A (Georgia) Tifton | 2002 | - | this paper | 1 | 1 | 0 |
| *P. ananatis* ^B^ | PANS 99-11 ^I^ | crab grass | U.S.A (Georgia) Tifton | 1999 | - | this paper | 1 | 1 | 0 |
| *P. ananatis* ^B^ | PNA 97-3 ^I^ | onion | - | 1997 | - | this paper | 1 | 1 | 1 |
| *P. ananatis* _B_ | PANS 99-09 ^I^ | prairie verbena | U.S.A (Georgia) Tifton | 1999 | - | this paper | 0 | 1 | 1 |
| *P. ananatis* ^B^ | PNA 98-7 ^I^ | onion | U.S.A (Georgia) Tifton | 1998 | - | this paper | 0 | 1 | 0 |
| *P. ananatis* ^B^ | PANS 01-10 ^I^ | thrips feces from peanut | U.S.A (Georgia) Tifton | 2001 | - | this paper | 0 | 0 | 0 |
| *P. ananatis* ^B^ | PANS 99-22 ^I^ | crab grass | U.S.A (Georgia) Tifton | 1999 | - | this paper | 0 | 0 | 0 |
| *P. ananatis* ^B^ | PANS 99-26 ^I^ | hyssop spurge | U.S.A (Georgia) Vidalia | 1999 | - | this paper | 0 | 0 | 0 |
| *P. ananatis* ^B^ | PANS 01-8 ^I^ | tobacco thrips | U.S.A (Georgia) Tifton | 2001 | - | this paper | 0 | 0 | 0 |
| *P. ananatis* ^B^ | PANS 01-09 ^I^ | thrips feces from peanut | U.S.A (Georgia) Tifton | 2001 | - | this paper | 0 | 0 | 0 |
| *P. ananatis* ^B^ | PNA 18-10S ^I^ | onion | U.S.A.(Georgia) Vidalia | 2018 | - | this paper | 0 | 0 | 0 |
| *P. ananatis* ^B^ | PNA 07-13 ^I^ | onion | - | 2007 | - | this paper | 0 | 1 | 0 |
| *P. ananatis* ^B^ | PNA 07-14 ^I^ | onion | - | 2007 | - | this paper | 0 | 1 | 0 |
| *P. ananatis* ^C^ | PNA 98-2 ^II^ | onion | U.S.A. (Georgia) - | 1998 | - | - | 1 | 1 | 1 |
| *P. stewartii* subsp. *indologenes* ^C^ | 0696-21 ^II^ | sudangrass | U.S.A. (California) Imperial Valley | 1996 | - | Azad et al. 2000 | 0 | 0 | 0 |
| *P. ananatis* ^D^ | LMG 2676 ^II^ | stem rust | U.S.A (-) - | 1954 | FJ611846 | Pon et al. 1954 | 0 | 0 | 0 |
| *P. ananatis* | LMG 2667 ^II^ | pineapple | U.S.A. (Hawaii) - | 1958 | - | - | 0 | 1 | 0 |
| *P. ananatis* | ATCC 35400 ^II^ | honey melons | Ecuador (-) - | 1981 | - | Wells et al. 1986 | 0 | 0 | 0 |
| *P. ananatis* ^C^ | Q12 ^II^ | onion | South Africa (-) - | - | - | - | 0 | 0 | 1 |
| *P. stewartii* subsp. *indologenes* ^D^ | LMG 2631 ^II^ | pearl millet | India (-) - | 1961 | KF482585 | - | 0 | 0 | 0 |
| *P. ananatis* ^D^ | Uruguay 37 ^II^ | eucalyptus | Uruguay (-) - | 2004 | - | - | 1 | 1 | 0 |
| *P. ananatis* ^D^ | BD 315 ^II^ | onion | U.S.A. (Georgia) - | - | AY579212 | Goszczynska et al. 2007 | 1 | 1 | 1 |
| *P. ananatis* | ICMP 10132 ^II^ | sugar cane | Brazil (-) - | 1991 | - | - | 0 | 0 | 0 |
| *P. allii* ^D^ | BD 309 ^II^ | onion | U.S.A. (-) - | 1994 | AY579210 | Goszczynska et al. 2011 | 0 | 0 | 1 |
| *P.allii* ^D^ | BD 377 ^II^ | onion | South Africa (-) - | 2004 | DQ512491 | Goszczynska et al. 2011 | 0 | 0 | 1 |
| *P. ananatis* ^D^ | BD 577 ^II^ | maize | South Africa (Mpumalanga) - | 2004 | DQ133547 | Goszczynska et al. 2007 | 0 | 0 | 0 |
| *P. stewartii* subsp. *indologenes* ^D^ | LC31 ^II^ | eucalyptus | South Africa (Mpumalanga) - | 2006 | - | - | 0 | 0 | 0 |
| *P. ananatis* ^D^ | LC 3 ^II^ | eucalyptus | South Africa (Mpumalanga) - | 2006 | - | - | 1 | 1 | 0 |
| *Pantoea* sp. ^E^ | 3095 ^II^ | - | U.S.A. (-) - | 2015 | - | - | 0 | 0 | 0 |
| *P. ananatis* ^D^ | BD 251 ^II^ | onion | South Africa (-) - | 2002 | - | - | 1 | 1 | 1 |
| *P. ananatis* ^D^ | BD 297 ^II^ | onion | South Africa (-) - | 2002 | - | - | 0 | 0 | 0 |
| *P. ananatis* ^D^ | BD 330 ^II^ | onion | South Africa (-) - | 2002 | - | - | 0 | 0 | 1 |
| *P. allii* ^D^ | BD 391 ^II^ | onion | South Africa (-) - | 2003 | - | - | 0 | 0 | 1 |
| *P. ananatis* ^D^ | BD 491 ^II^ | maize | South Africa (-) - | 2005 | - | - | 1 | 1 | 0 |
| *P. ananatis* ^D^ | BD 570 ^II^ | maize | South Africa (-) - | 2005 | - | - | 0 | 0 | 0 |
| *P. ananatis* ^D^ | BD 647 ^II^ | maize | South Africa (-) - | 2005 | DQ195525 | Goszczynska et al. 2007 | 1 | 1 | 1 |
| *P. ananatis* ^D^ | SUPP2113 ^II^ | rice | Japan (-) - | 2004 | - | Kido et al. 2010 | 1 | 1 | 0 |
| *P. ananatis* ^D^ | MBB 35+ ^II^ | eucalyptus | South Africa (KwaZulu-Natal) Mtunzini | - | - | - | 0 | 0 | 0 |
| *P. dispersa* ^D^ | TMA 7 ^II^ | eucalyptus | Thailand (-) - | - | - | - | 0 | 0 | 0 |
| *P. ananatis* | LMG 20104 ^II^ | eucalyptus | South Africa (KwaZulu-Natal) Harding | 2001 | - | Coutinho et al. 2002 | 0 | 0 | 0 |
| *P. ananatis* | 0197-28 ^II^ | sudangrass | U.S.A. (California) Imperial Valley | 1996 | - | Azad et al. 2000 | 0 | 0 | 0 |
| *P. ananatis* | LMG 2666 ^II^ | pineapple | U.S.A. (Hawaii) - | 1957 | - | - | 1 | 1 | 0 |
| *P. ananatis* ^C^ | CTB1135 ^II^ | rice | Japan (Chūgoku) Tottori | 1995 | - | Kido et al. 2008 | 1 | 0 | 0 |
| *P. ananatis* | LMG 2101 ^II^ | rice | India (-) - | 2001 | - | - | 0 | 0 | 0 |
| *P. ananatis* ^D^ | DAR76141 ^II^ | rice | Australia (New South Wales) Whitton | 2004 | - | Cother et al. 2004 | 1 | 1 | 0 |
| *P. agglomerans* ^D^ | LMG 2596 ^II^ | onion | South Africa (Western Cape) Little Karoo | 1977 | EF988816 | Hattingh & Walters 1981 | 1 | 0 | 1 |
| *P. ananatis* ^D^ | BD 301 ^II^ | onion | U.S.A. (Georgia) Tifton | - | AY579209 | Goszczynska et al. 2006 | 1 | 1 | 1 |
| *P. ananatis* ^D^ | BD 315 ^II^ | onion | U.S.A. (Georgia) - | - | AY579212 | Goszczynska et al. 2006 | 1 | 1 | 1 |
| *P. ananatis* ^D^ | BD 435 ^II^ | maize | South Africa (Mpumalanga) - | 2004 | AY898642 | Goszczynska et al. 2007 | 0 | 0 | 0 |
| *P. ananatis* ^D^ | BD 543 | maize | South Africa (Northwest) - | 2004 | DQ133545 | Goszczynska et al. 2007 | 0 | 0 | 0 |
| *P. ananatis* ^D^ | BD 561 ^II^ | maize | South Africa (Northwest) - | 2004 | DQ133546 | Goszczynska et al. 2007 | 0 | 0 | 0 |
| *P. ananatis* ^D^ | BD 588 ^II^ | maize | South Africa (Mpumalanga) - | 2004 | DQ133548 | Goszczynska et al. 2007 | 1 | 1 | 0 |
| *P. ananatis* ^D^ | BD 647 ^II^ | maize | South Africa (Mpumalanga) - | 2004 | DQ195525 | Goszczynska et al. 2007 | 1 | 1 | 0 |
| *P. allii* ^D^ | BD 304 ^II^ | onion | South Africa (-) - | 2002 | - | - | 0 | 1 | 0 |
| *P. ananatis* ^D^ | BD 546 ^II^ | maize | South Africa (-) - | 2005 | - | - | 1 | 1 | 0 |
| *P. vagans* ^C^ | LMG 24196 ^II^ | eucalyptus | Argentina (-) - | - | EF988758 | Brady et al. 2009 | 1 | 0 | 0 |
| *P. dispersa* ^D^ | TMA 3 ^II^ | eucalyptus | Thailand (-) - | - | - | - | 0 | 0 | 0 |
| *P. ananatis* ^C^ | Mmir 7+ ^II^ | capsid bug | South Africa (-) - | - | - | - | 0 | 0 | 0 |
| *P. ananatis* ^C^ | Mmir 8+ ^II^ | capsid bug | South Africa (-) - | - | - | - | 0 | 0 | 0 |
| *P. ananatis* ^C^ | Mmir 10+ ^II^ | capsid bug | South Africa (-) - | - | - | - | 1 | 1 | 0 |
| *P. eucalypti* ^A^ | LMG 24197 ^T, II^ | eucalyptus | Uruguay (-) - | - | VHJB00000000 | Brady et al. 2009 | 0 | 0 | 0 |
| *P. eucalypti* ^D^ | LMG 24198 ^II^ | eucalyptus | Uruguay (-) - | - | EF988763 | Brady et al. 2009 | 0 | 0 | 0 |
| *P. vagans* ^C^ | BCC0079 ^II^ | eucalyptus | Uruguay (-) - | - | - | Brady et al. 2008 | 0 | 0 | 0 |
| *P. ananatis* ^C^ | BCC0083 ^II^ | onion | U.S.A. (-) - | - | - | Brady et al. 2007 | 1 | 1 | 1 |
| *P. vagans* ^A^ | LMG 24199 ^T, II^ | eucalyptus | Uganda (-) - | - | CP038853 - CP038855 | Brady et al. 2008 | 0 | 0 | 0 |
| *P. deleyi* ^A^ | LMG 24200 ^T, II^ | eucalyptus | Uganda (-) - | - | MIPO00000000 | Brady et al. 2008 | 0 | 0 | 0 |
| *P. stewartii* subsp*. indologenes* ^C^ | BCC0118 ^II^ | eucalyptus | South Africa (KwaZulu-Natal) Futululu | - | - | Brady et al. 2008 | 0 | 0 | 0 |
| *P. stewartii* subsp. *indologenes* ^E^ | MKB 0035 ^II^ | eucalyptus | South Africa (KwaZulu-Natal) Futululu | - | - | - | 0 | 0 | 0 |
| *P. stewartii* subsp. *indologenes* ^C^ | LMG 2671 ^II^ | pineapple | U.S.A. (Hawaii) - | 1948 | EF988826 | Brady et al. 2008 | 0 | 0 | 0 |
| *P. agglomerans* ^C^ | LMG 1286 ^II^ | human wound | Zimbabwe (-) - | 1956 | NR_116751 | Rezzonico et al. 2009 | 0 | 0 | 0 |
| *P. stewartii* subsp. *indologenes* ^A^ | LMG 2632 ^T, II^ | foxtail millet | India (-) - | 1960 | JPKO00000000 | Brady et al. 2008 | 0 | 0 | 0 |
| *P. agglomerans* | BD 176 ^II^ | onion | South Africa (-) - | - | - | - | 0 | 0 | 0 |
| *P. agglomerans* | BD 109 ^II^ | onion | South Africa (-) - | - | - | - | 0 | 0 | 1 |
| *P. ananatis* ^D^ | LMG 2680 ^II^ | stem rust | Hungary (-) - | 1956 | - | Mergaert et al. 1993 | 0 | 0 | 0 |
| *P. ananatis* ^D^ | LMG 2807 ^II^ | orchid | Brazil (-) - | 1965 | - | Gehring et al. 2014 | 1 | 0 | 0 |
| *P. dispersa* ^C^ | LMG 2604 ^II^ | wild rose | Netherlands (-) - | 1969 | EF988819 | Brady et al. 2008 | 0 | 0 | 0 |
| *P. agglomerans* ^C^ | LMG 2660 ^II^ | japanese wistaria | Japan (-) - | 1970 | Z96083 | Hauben et al. 1998 | 0 | 0 | 0 |
| *P. dispersa* ^C^ | LMG 2749 ^II^ | human wound | - (-) - | 1979 | EF988833 | Brady et al. 2008 | 0 | 0 | 0 |
| *P. agglomerans* ^C^ | LMG 2565 ^II^ | oat | Canada (-) - | 1979 | Z96082 | Hauben et al. 1998 | 0 | 0 | 0 |
| *P. agglomerans* | LMG 2570 ^II^ | mountain-ash | U.S.A. (-) - | 1980 | - | Buttimer et al. 2017 | 0 | 0 | 0 |
| *P. dispersa* ^C^ | LMG 2603 ^T, II^ | soil | Japan (-) - | 1979 | NR_043883 | Chang et al. 2018 | 0 | 0 | 0 |
| *P. stewartii* subsp. *stewartia* ^C^ | LMG 2715 ^T, II^ | maize | U.S.A. (-) - | - | Z96080 | Hauben et al. 1998 | 0 | 0 | 0 |
| *P. agglomerans* ^E^ | Uruguay 41 ^II^ | eucalyptus | Uruguay (-) - | 2004 | - | - | 0 | 0 | 0 |
| *P. ananatis* ^D^ | BD 335 ^II^ | onion | South Africa (-) - | - | - | - | 0 | 0 | 1 |
| *P. agglomerans* pv*. gypsophilae* | LMG 2553 ^II^ | baby's-breath | U.S.A. | 1946 | EF988810 | Brady et al. 2008 | 0 | 0 | 0 |
| *P. allii* ^A^ | LMG 24248 ^T, II^ | onion | South Africa (-) - | 2004 | MLFE00000000 | Brady et al. 2011 | 0 | 0 | 1 |
| *P. ananatis* ^A^ | BD 442 ^II^ | maize | South Africa (Mpumalanga) - | 2004 | JMJL00000000 | Goszczynska et al. 2007 | 0 | 0 | 0 |
| *P. ananatis* | BD 640 ^II^ | maize | South Africa (Mpumalanga) - | 2004 | DQ195524 | Goszczynska et al. 2007 | 0 | 0 | 0 |
| *P. wallisii* ^A^ | LMG 26277 ^T, II^ | eucalyptus | South Africa (White River) - | 2006 | MLFS00000000 | Brady et al. 2012 | 0 | 0 | 0 |
| *P. eucalypti* | LC53 ^II^ | eucalyptus | South Africa (White River) - | 2006 | - | - | 0 | 0 | 0 |
| *P. agglomerans* ^C^ | LMG 2941 ^II^ | crab apple | - (-) - | 1979 | FJ611837 | Rezzonico et al. 2009 | 0 | 0 | 0 |
| *P. anthophila* ^D^ | LMG 2558 ^T, II^ | garden balsam | India (-) - | 1981 | VHIZ00000000 | Brady et al. 2009 | 0 | 0 | 0 |
| *P. agglomerans* pv*. betae* ^C^ | PAB4188 ^II^ | beet | U.S.A. (New York) Geneva | | - | Burr et al. 1991 | 0 | 0 | 0 |
| *P. dispersa* ^D^ | LMG 2605 ^II^ | cowpea | Tanzania (-) - | 1965 | FJ611866 | Rezzonico et al. 2009 | 0 | 0 | 0 |
| *P. agglomerans* ^D^ | LMG 2554 ^II^ | runner bean | U.K. (-) - | 1981 | EF988811 | Brady et al. 2008 | 0 | 0 | 0 |
| *P. agglomerans* ^D^ | LMG 2572 ^II^ | wheat | Canada (-) - | - | EF988815 | Brady et al. 2008 | 0 | 0 | 0 |
| *P. eucalypti* ^D^ | Uruguay 17 ^II^ | eucalyptus | Uruguay (-) - | 2004 | - | - | 0 | 0 | 0 |
| *P. eucalypti* ^D^ | 31b g ^II^ | eucalyptus | Uruguay (-) - | 2004 | - | - | 0 | 0 | 0 |
| *P. agglomerans* ^D^ | NCCP222 ^II^ | soil | Pakistan (-) - | 2010 | - | - | 0 | 0 | 0 |
| *P. agglomerans* ^D^ | NCCP222 ^II^ | soil | Pakistan (-) - | 2010 | - | - | 0 | 0 | 0 |
| *P. rodasii* ^D^ | BD943 ^II^ | eucalyptus | Colombia (-) - | 2005 | MLFP00000000 | Brady et al. 2012 | 0 | 0 | 0 |
| *P. wallisii* ^D^ | BD946 ^II^ | eucalyptus | South Africa (-) - | 2006 | MLFS00000000 | Brady et al. 2012 | 0 | 0 | 0 |
| *P. rwandensis ­*^D^ | BD944 ^II^ | eucalyptus | Rwanda (-) - | 2005 | MLFR00000000 | Brady et al. 2012 | 0 | 0 | 0 |
| *P. beijingensis* ^D^ | BCC1348 ^II^ | oyster mushroom | China (-) - | 2014 | JMEE00000000 | Liu et al. 2013 | 0 | 0 | 0 |
| *P. beijingensis* ^D^ | BCC1349 ^II^ | oyster mushroom | China (-) - | 2014 | KC846071 | Liu et al. 2013 | 0 | 0 | 0 |
| *P. pleuroti* ^D^ | BCC1352 ^II^ | oyster mushroom | China (-) - | 2014 | KJ654341 | Ma et al. 2016 | 0 | 0 | 0 |
| *P. pleuroti* ^D^ | BCC1353 ^II^ | oyster mushroom | China (-) - | 2014 | KJ654341 | Ma et al. 2016 | 0 | 0 | 0 |
| *P. ananatis* ^A^ | AJ13355 ^II^ | soil | Japan (-) - | - | NC_017531 - NC_017533 | Hara et al. 2012 | 0 | 0 | 0 |
| *P. eucalypti* ^D^ | BD 300 ^II^ | onion | South Africa (-) - | 2002 | - | - | 0 | 0 | 1 |
| *P. ananatis* ^D^ | BD 307 ^II^ | onion | South Africa (-) - | 2002 | - | - | 0 | 0 | 1 |
| *P. agglomerans* ^D^ | BD 378 ^II^ | onion | South Africa (-) - | 2003 | - | - | 0 | 0 | 1 |
| *P. allii* ^D^ | BD 379 ^II^ | onion | South Africa (-) - | 2003 | - | - | 0 | 0 | 1 |
| *P. allii* ^D^ | BD 380 ^II^ | onion | South Africa (-) - | 2003 | GU458415 | Goszczynska et al. 2011 | 1 | 0 | 1 |
| *P. allii* ^D^ | BD 381 ^II^ | onion | South Africa (-) - | 2003 | GU458416 | Goszczynska et al. 2011 | 1 | 0 | 1 |
| *P. allii* ^D^ | BD 382 ^II^ | onion | South Africa (-) - | 2003 | - | - | 1 | 0 | 0 |
| *P. allii* ^D^ | BD 383 ^II^ | onion | South Africa (-) - | 2003 | GU458417 | Goszczynska et al. 2011 | 1 | 0 | 1 |
| *P. allii* ^D^ | BD 384 ^II^ | onion | South Africa (-) - | 2003 | - | - | 1 | 0 | 1 |
| *P. allii* ^D^ | BD 385 ^II^ | onion | South Africa (-) - | 2003 | - | - | 1 | 0 | 1 |
| *P. allii* ^D^ | BD 386 ^II^ | onion | South Africa (-) - | 2003 | - | - | 1 | 0 | 1 |
| *P. allii* ^D^ | BD 387 ^II^ | onion | South Africa (-) - | 2003 | - | - | 1 | 0 | 1 |
| *P. allii* ^D^ | BD 388 ^II^ | onion | South Africa (-) - | 2003 | - | - | 1 | 0 | 1 |
| *P. allii* ^D^ | BD 389 ^II^ | onion | South Africa (-) - | 2003 | - | - | 1 | 0 | 1 |
| *P. conspicua* ^D^ | BD 479 ^II^ | maize | South Africa (-) - | 2005 | - | - | 0 | 1 | 0 |
| *P. anthophila* ^D^ | BD 589 ^II^ | maize | South Africa (-) - | 2005 | - | - | 0 | 0 | 0 |
| *P. anthophila* ^D^ | BD 590 ^II^ | maize | South Africa (-) - | 2005 | - | - | 0 | 0 | 0 |
| *P. eucrina* ^D^ | BD 591 ^II^ | maize | South Africa (-) - | 2005 | - | - | 0 | 0 | 0 |
| *P. eucrina* ^D^ | BD 592 ^II^ | maize | South Africa (-) - | 2005 | - | - | 0 | 0 | 0 |
| *P. stewartii* ^D^ | BD 641 ^II^ | maize | South Africa (-) - | 2005 | - | - | 0 | 0 | 0 |
| *P. anthophila* ^D^ | BD 644 ^II^ | maize | South Africa (-) - | 2005 | - | - | 0 | 0 | 0 |
| *P. anthophila* ^D^ | BD 645 ^II^ | maize | South Africa (-) - | 2005 | - | - | 0 | 0 | 0 |
| *P. agglomerans* ^D^ | BD 653 ^II^ | maize | South Africa (-) - | 2005 | - | - | 0 | 0 | 0 |
| *P. eucrina* ^D^ | BD 680 ^II^ | watermelon | South Africa (-) - | 2005 | - | - | 0 | 0 | 0 |
| *P. agglomerans* ^D^ | BD 689 ^II^ | onion | South Africa (-) - | 2005 | - | - | 0 | 0 | 1 |
| *Pantoea* sp*.* ^D^ | YR343 ^II^ | eastern cottonwood | - (-) - | 2012 | AKIT00000000 | Brown et al. 2012 | 0 | 0 | 0 |
| *P. ananatis* ^D^ | ICMP10132 ^II^ | maize | Brazil (-) - | 1989 | - | - | 0 | 0 | 0 |
| *P. stewartii* subsp. *indologenes* ^D^ | ICMP 12183 ^II^ | maize | Brazil (-) - | 1991 | - | - | 0 | 0 | 0 |
| *P. vagans* ^D^ | MP7 ^II^ | termites | South Africa (-) - | 2012 | JPKP00000000 | Palmer et al. 2016 | 0 | 0 | 0 |
| *P. agglomerans* ^D^ | MAI 6000 ^III^ | onion | Uruguay (Canelones) - | 2015 |  |  | 0 | 0 | 1 |
| *P. agglomerans* ^D^ | MAI 6001 ^III^ | onion | Uruguay (Canelones) - | 2015 |  |  | 0 | 0 | 1 |
| *P. agglomerans* ^D^ | MAI 6002 ^III^ | onion | Uruguay (Canelones) - | 2015 |  |  | 0 | 0 | 1 |
| *P. eucalypti* ^D^ | MAI 6003 ^III^ | onion | Uruguay (Canelones) - | 2015 |  |  | 0 | 0 | 0 |
| *P. allii* ^D^ | MAI 6004 ^III^ | onion | Uruguay (Canelones) - | 2018 |  |  | 0 | 0 | 1 |
| *P. agglomerans* ^D^ | MAI 6005 ^III^ | onion | Uruguay (Canelones) - | 2018 |  |  | 0 | 0 | 1 |
| *P. allii* ^D^ | MAI 6006 ^III^ | onion | Uruguay (Canelones) - | 2018 |  |  | 0 | 0 | 1 |
| *P. allii* ^D^ | MAI 6007 ^III^ | onion | Uruguay (Canelones) - | 2018 |  |  | 0 | 0 | 0 |
| *P. eucalypti* ^D^ | MAI 6008 ^III^ | onion | Uruguay (Canelones) - | 2018 |  |  | 0 | 0 | 0 |
| *P. eucalypti* ^D^ | MAI 6009 ^III^ | onion | Uruguay (Canelones) - | 2018 |  |  | 0 | 0 | 1 |
| *P. agglomerans* ^D^ | MAI 6010 ^III^ | onion | Uruguay (Canelones) - | 2018 |  |  | 0 | 0 | 1 |
| *P. eucalypti* ^D^ | MAI 6011 ^III^ | onion | Uruguay (Canelones) - | 2018 |  |  | 0 | 0 | 1 |
| *P. agglomerans* ^D^ | MAI 6012 ^III^ | onion | Uruguay (Canelones) - | 2018 |  |  | 0 | 0 | 1 |
| *P. eucalypti* ^D^ | MAI 6013 ^III^ | onion | Uruguay (Canelones) - | 2018 |  |  | 0 | 0 | 1 |
| *P. eucalypti* ^D^ | MAI 6014 ^III^ | onion | Uruguay (Canelones) - | 2018 |  |  | 0 | 0 | 1 |
| *P. eucalypti* ^D^ | MAI 6015 ^III^ | onion | Uruguay (Canelones) - | 2018 |  |  | 0 | 0 | 1 |
| *P. eucalypti* ^D^ | MAI 6016 ^III^ | onion | Uruguay (Canelones) - | 2018 |  |  | 0 | 0 | 1 |
| *P. agglomerans* ^D^ | MAI 6017 ^III^ | onion | Uruguay (Canelones) - | 2018 |  |  | 0 | 0 | 1 |
| *P. agglomerans* ^D^ | MAI 6018 ^III^ | onion | Uruguay (Canelones) - | 2018 |  |  | 0 | 0 | 1 |
| *P. eucalypti* ^D^ | MAI 6019 ^III^ | onion | Uruguay (Canelones) - | 2018 |  |  | 0 | 0 | 1 |
| *Pantoea* sp. ^D^ | MAI 6020 ^III^ | onion | Uruguay (Canelones) - | 2018 |  |  | 0 | 0 | 1 |
| *P. eucalypti* ^D^ | MAI 6021 ^III^ | onion | Uruguay (Canelones) - | 2018 |  |  | 0 | 0 | 0 |
| *P. allii* ^D^ | MAI 6022 ^III^ | onion | Uruguay (Canelones) - | 2018 |  |  | 0 | 0 | 1 |
| *P. agglomerans* ^D^ | MAI 6023 ^III^ | onion | Uruguay (Canelones) - | 2018 |  |  | 0 | 0 | 1 |
| *P. eucalypti* ^D^ | MAI 6024 ^III^ | onion | Uruguay (Canelones) - | 2018 |  |  | 0 | 0 | 1 |
| *P. agglomerans* | MAI 6025 ^III^ | onion | Uruguay (Canelones) - | 2018 |  |  | 0 | 0 | 1 |
| *P. eucalypti* ^D^ | MAI 6026 ^III^ | onion | Uruguay (Salto) - | 2018 |  |  | 0 | 0 | 1 |
| *P. eucalypti* ^D^ | MAI 6027 ^III^ | onion | Uruguay (Salto) - | 2018 |  |  | 0 | 0 | 0 |
| *P. eucalypti* ^D^ | MAI 6028 ^III^ | onion | Uruguay (Salto) - | 2018 |  |  | 0 | 0 | 1 |
| *P. eucalypti* ^D^ | MAI 6029 ^III^ | onion | Uruguay (Salto) - | 2018 |  |  | 0 | 0 | 1 |
| *P. eucalypti* ^D^ | MAI 6030 ^III^ | onion | Uruguay (Salto) - | 2018 |  |  | 0 | 0 | 1 |
| *P. eucalypti* ^D^ | MAI 6031 ^III^ | onion | Uruguay (Salto) - | 2018 |  |  | 0 | 0 | 1 |
| *P. ananatis* ^D^ | MAI 6032 ^III^ | onion | Uruguay (Salto) - | 2018 |  |  | 1 | 1 | 1 |
| *P. eucalypti* ^D^ | MAI 6033 ^III^ | onion | Uruguay (Salto) - | 2018 |  |  | 0 | 0 | 1 |
| *P. eucalypti* ^D^ | MAI 6034 ^III^ | onion | Uruguay (Salto) - | 2018 |  |  | 0 | 0 | 1 |
| *P. eucalypti* ^D^ | MAI 6035 ^III^ | onion | Uruguay (Salto) - | 2018 |  |  | 0 | 0 | 1 |
| *P. eucalypti* ^D^ | MAI 6036 ^III^ | onion | Uruguay (Salto) - | 2018 |  |  | 1 | 0 | 1 |
| *P. eucalypti* ^D^ | MAI 6037 ^III^ | onion | Uruguay (Salto) - | 2018 |  |  | 0 | 0 | 0 |
| *P. eucalypti* ^D^ | MAI 6038 ^III^ | onion | Uruguay (Salto) - | 2018 |  |  | 0 | 0 | 1 |
| *P. ananatis* ^D^ | MAI 6039 ^III^ | onion | Uruguay (Salto) - | 2018 |  |  | 1 | 1 | 1 |
| *P. eucalypti* ^D^ | MAI 6040 ^III^ | onion | Uruguay (Salto) - | 2018 |  |  | 0 | 0 | 1 |
| *P. eucalypti* ^D^ | MAI 6041 ^III^ | onion | Uruguay (Salto) - | 2018 |  |  | 0 | 0 | 1 |
| *P. eucalypti* ^D^ | MAI 6042 ^III^ | onion | Uruguay (Salto) - | 2018 |  |  | 0 | 0 | 1 |
| *P. eucalypti* ^D^ | MAI 6043 ^III^ | onion | Uruguay (Salto) - | 2019 |  |  | 0 | 0 | 0 |
| *P. eucalypti* ^D^ | MAI 6044 ^III^ | onion | Uruguay (Salto) - | 2019 |  |  | 0 | 0 | 1 |
| *P. agglomerans* ^D^ | MAI 6045 ^III^ | onion | Uruguay (Salto) - | 2019 |  |  | 1 | 0 | 1 |
| *P. eucalypti* ^D^ | MAI 6046 ^III^ | onion | Uruguay (Salto) - | 2019 |  |  | 0 | 0 | 1 |
| *P. eucalypti* ^D^ | MAI 6047 ^III^ | onion | Uruguay (Salto) - | 2019 |  |  | 0 | 0 | 1 |
| *P. eucalypti* ^D^ | MAI 6048 ^III^ | onion | Uruguay (Salto) - | 2019 |  |  | 0 | 0 | 0 |
| *Pantoea* sp. ^D^ | MAI 6049 ^III^ | onion | Uruguay (Salto) - | 2019 |  |  | 0 | 0 | 1 |
| *P. vagans* ^D^ | MAI 6050 ^III^ | onion | Uruguay (Salto) - | 2019 |  |  | 0 | 0 | 0 |
| *P. eucalypti* ^D^ | MAI 6051 ^III^ | onion | Uruguay (Salto) - | 2019 |  |  | 0 | 0 | 1 |
| *P. eucalypti* ^D^ | MAI 6052 ^III^ | onion | Uruguay (Salto) - | 2019 |  |  | 0 | 0 | 1 |
| *Pantoea* sp. ^D^ | MAI 6053 ^III^ | onion | Uruguay (Salto) - | 2019 |  |  | 0 | 0 | 1 |

Sequence identification method: ^A^ Whole genome sequencing (WGS), ^B^ Species specific primers, ^C^ Amplified fragment length polymorphism (AFLP), ^D^ Multi Locus Sequence Analysis (MLSA), ^E^ 16S rRNA gene sequence. Bacterial culture collection: ^I^ UGA-CPES, ^II^ UP-BCC, ^III^ UR-MAI. RSN = Red scale necrosis ([1] clearly defined lesion created by bacteria on detached red onion scale 3 DPI; [0] no lesion created by bacteria 3 DPI). HiVir2p_F/R = High Virulence cluster PCR assay result ([1] obvious visible amplicon 857 bp; [0] no visible amplicon or not interpretable). alt1p_F/R = allicin tolerance cluster PCR assay result ([1] obvious visible amplicon 414 bp; [0] no visible amplicon or not interpretable)

**Supplementary Table 2.** Z-proportional test comparing group totals from Table 2. *Pnan* = *Pantoea ananatis*, *Pn* spp. = *Pantoea* spp. Onion = bacteria originally isolated from diseased onion tissue. Non-onion = bacteria isolate from any sources other than diseased onion tissue. RSN = Red scale necrosis (+ clearly defined lesion created by bacteria on detached red onion scale 3 DPI; – no lesion created by bacteria 3 DPI). *alt* = allicin tolerance cluster PCR assay result (+ obvious visible amplicon 414bp; - no visible amplicon or not interpretable).

|  | **Z-proportions test** | | | |
| --- | --- | --- | --- | --- |
| **Comparison** | RSN+ *alt*+ | RSN- *alt*+ | RSN+ *alt-* | RSN- *alt*- |
| *Pnan* onion vs. *Pnan* non-onion | Z=4.062, P<0.00001 | Z=3.103,  P = 0.0019 | Z=-3.55, P=0.00038 | Z=0.416, P=0.68 |
| Pn spp onion vs. Pn spp non-onion | Z=3.345, P=0.0008 | Z=7.244, P<0.00001 | Z=-0.18, P=0.86 | Z=-8.93, P<0.00001 |
| Pnan onion vs. Pn spp onion | Z=4.103, P<0.00001) | Z=-4.67, P<0.00001 | Z=0.96, P=0.34 | Z=0.74, P=0.46 |
| Pnan non-onion vs. Pn spp non-onion | Z=3.193, P=0.0014 | Z=-0.066, P=0.94 | Z=4.24, P<0.00001 | Z=-5.41, P<0.00001 |
| Pnan onion vs. Pn spp non-onion | Z=6.32,  P < 0.00001 | Z= 2.98,  P < 0.003 | Z=0.73,  P =0.47 | Z=-7.74 P<0.00001 |
| Pnan non-onion vs. Pn spp onion | Z=-0.2,  P =0.84 | Z=-7.45,  P<0.0001 | Z=4.81, P<0.00001 | Z=4.19, P<0.00001 |
